# Physiological architecture and evolutionary origins of cellular adaptability

**DOI:** 10.64898/2026.04.10.717775

**Authors:** Annisa Dea, Yongqing Lan, Benjamin A. Doran, Asif Ali, Maya G. Igarashi, Lucas Dyer, Valeryia Aksianiuk, David Pincus, Arjun S. Raman

**Affiliations:** Department of Molecular Genetics and Cell Biology, University of Chicago, Chicago, IL; Duchossois Family Institute, University of Chicago, Chicago, IL; Department of Pathology, University of Chicago, Chicago, IL; Pritzker School of Molecular Engineering, University of Chicago, Chicago, IL; Institute for Biophysical Dynamics, University of Chicago, Chicago, IL; Center for Physics of Evolving Systems, University of Chicago, Chicago, IL

## Abstract

The capacity to adapt to complex environments (‘adaptability’) is a defining property of cells. Yet, its relationship across single-cell physiology, population-level responses, and evolutionary timescales remains unclear. We performed single-cell transcriptional profiling of budding yeast across 20 complex environments before and after long-term selection for increased osmotolerance. In ancestral populations, transcriptional responses organized into a reproducible hierarchy where adaptation to certain cues took precedence over others. This hierarchy mechanistically originated within individual cells through contingent regulation of translation initiation. Evolution for >3,000 generations under osmotic stress increased osmotolerance while reordering this hierarchy, selectively deprioritizing osmotic stress as an organizing axis of adaptation. The derived strain exhibited impaired integration of stress responses, defective translational coordination, and reduced fitness outside the selected condition. Together, these findings illustrate that cellular adaptability is an evolved architecture whose form is set by the history of selection.

**One Sentence Summary:** Cells adapt to complex environments via a prioritization hierarchy; this hierarchy is shaped by the frequency and magnitude of environmental fluctuations in their evolutionary history.

## Introduction

Living systems persist in environments that can change unpredictably. Thus, survival depends on the capacity to adapt (e.g. ‘adaptability’)—a systems-level property emerging from interacting molecular components and regulatory layers expressed as a coherent, organized phenotypic logic [Anderson, 1972, Jacob and Monod, 1961, Balch et al., 2008, Morimoto, 2011, Lindquist, 1986]. Previous work on cellular adaptation has mostly examined the effects of single environmental perturbations. From this approach a useful organizing picture has emerged in which cells harbor a shared, global environmental stress response layered with condition-specific transcriptional programs [Gasch et al., 2000, Gasch and Werner-Washburne, 2002, Berry and Gasch, 2008, Brauer et al., 2008, Capaldi et al., 2008]. However, cells rarely experience stresses in isolation. Natural environments vary along multiple axes, and combined perturbations can interact, imposing competing or synergistic physiological demands. These observations raise a key question: in complex environments, is adaptability governed by a stereotyped organizing structure, or does it fragment into context-specific programs that must be re-interpreted for each distinct environment?

Addressing this question has been challenging as the space of possible environments is combinatorially expansive, and responses to combined perturbations can be non-additive reflecting nonlinear interactions across regulatory layers [McClean et al., 2007, Hao and O’shea, 2012, Vidal et al., 2013]. Moreover, similar physiological challenges can be resolved through multiple molecular routes, while distinct stresses can converge on shared bottlenecks—obscuring common structure when analyses focus on individual genes or individual conditions. Thus, it is difficult to determine whether responses across complex environments respect an underlying molecular logic or are fundamentally high-dimensional and idiosyncratic.

Here, we test whether responses across complex environments exhibit shared structure. We chose *S. cerevisiae* as our model system because its molecular biology is extensively mapped, it can be robustly cultured across diverse conditions, and it is supported by a mature genetic toolkit for perturbation and manipulation. We profiled yeast single-cell transcriptomes across 20 environments, varying carbon source, nutrient richness, osmolarity, temperature, and reactive oxygen species—thereby sampling combinations of well-studied individual stresses [Gasch et al., 2000, Brewster and Gustin, 2014, Anckar and Sistonen, 2011, Ma, 2013, Kourtis and Tavernarakis, 2011, de Nadal et al., 2011, Dea and Pincus, 2024]. Rather than analyzing each condition in isolation, we treated cellular transcriptomes measured across all environments as a single ensemble and interrogated whether there were statistical patterns of transcriptional variation that organize adaptive responses. Our results revealed a hierarchical structure of adaptation where variation in transcriptional responses was ordered by sensitivity to carbon source first, then nutrient richness, osmolarity, temperature, and finally reactive oxygen species. This hierarchy suggested that some environmental features limit the range of responses available to others, effectively gating how cells respond in combined environments. We experimentally validated these predictions within individual cell populations. Further, we found that this prioritization logic manifested mechanistically within single cells, emerging from differential regulation of translation initiation and synthesis of ribosomal proteins. Finally, we evolved yeast under constant osmotic stress (>3,000 generations; ∼9 months of continuous culture) and re-profiled evolved populations at single-cell resolution across the same panel of 20 environments. Long-term evolution reordered this hierarchy, selectively deprioritizing osmotic stress and impairing the integration of combined stresses despite increased fitness in the selected condition.

Together, these results demonstrate a hierarchical logic by which cells adapt to complex environments—a finding that refines the well-established model of the ‘environmental stress response’ (ESR) [Gasch et al., 2000, Gasch and Werner-Washburne, 2002]. We find this logic is encoded at the scale of individual cells within a population via differential translation initiation [Costa-Mattioli and Walter, 2020, Iserman et al., 2020]. Importantly, our results show that this logic is not fixed; rather it is shaped by the history of selection [Rutherford and Lindquist, 1998, Queitsch et al., 2002, Cowen and Lindquist, 2005, Jarosz and Lindquist, 2010]. Thus, our results show that the architecture of adaptability is itself an evolved trait, capable of being sculpted by the evolutionary history of the species.

## Results

### A statistical hierarchy orders adaptation by environmental cues

We grew haploid *S. cerevisiae* cells in different environments in which we varied carbon source (glucose or ethanol + glycerol), media composition (yeast extract with peptone (YP) or synthetic complete (SC)), temperature (30°C or 37°C), osmolarity (isotonic, KCl, or sorbitol), and reactive oxygen species (ROS) (none, H_2_O_2_, or diamide) (**Fig. 1A**). We selected 20 combinations of these environmental cues that resulted in a broad range of observed growth rates (**Fig. 1B; fig. S1A, B**). We grew cultures to maximal logarithmic phase growth rate in each of these environments to allow cells to adapt physiologically (**Methods**). We then harvested cells for single cell transcriptional profiling (scRNA-seq); in each environment, we sequenced ≥ 1000 cells with ≥ 500 genes expressed (**Fig. 1C**, **fig. S2**, Table S1) (**Methods**) [Gasch et al., 2017, Nadal-Ribelles et al., 2019, Jariani et al., 2020]. Uniform manifold approximation and projection (UMAP) revealed separation of cells into clusters based on the environment in which they were grown (**Fig. 1D**). Cells grown in ethanol + glycerol were fully separated from cells grown in glucose, and cells grown in hyperosmotic environments largely separated from isotonic cells, but variation in other environmental categories was less apparent in the UMAP visualization (**fig. S3**).

**Fig. 1:**
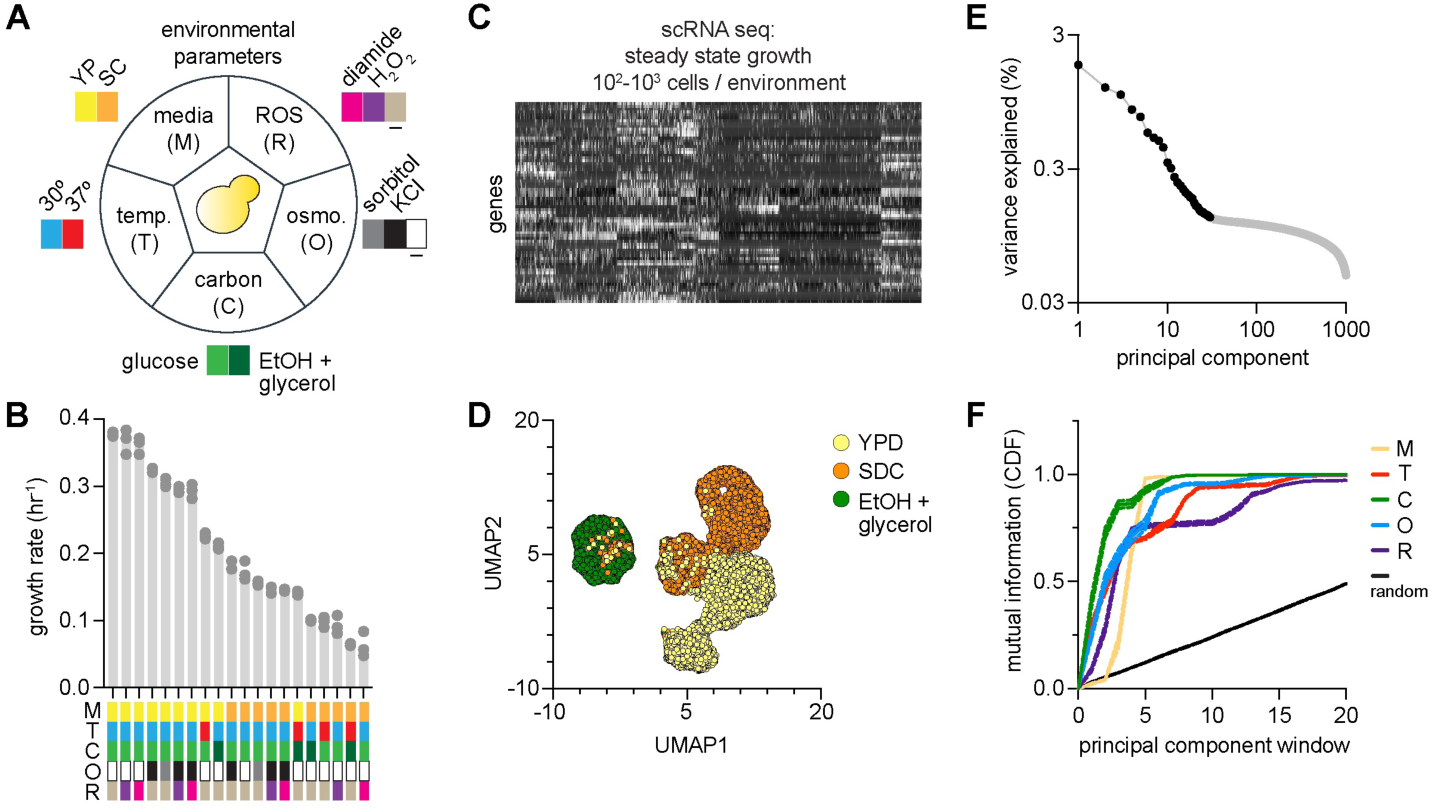
A statistical hierarchy orders cellular responses to complex environments. **A.** Set of environmental cues used to define complex environments in which *S. cerevisiae* was grown (YP: yeast extract + peptone; SC: synthetic complete; ROS: reactive oxygen species). **B.** Growth rate of population (y-axis) versus composition of complex environments (x-axis). **C, D.** Single-cell transcriptional profiling (scRNA-seq) performed across all cell populations (panel C). UMAP clustering illustrates separation of cell populations by specific environmental cues (panel D). **E.** Eigenspectrum of data shown in panel C. Black dots highlight principal components (x-axis) up to which linear scaling with percent variance explained (y-axis) was observed. **F.** Information shared (y-axis) between statistical structure of transcriptional variation (x-axis) and environmental cues (color key). Mutual information (MI) plotted as cumulative distribution (CDF). ‘Random’ reflects null expectation of MI.

We next interrogated the statistical structure of transcriptional variation across all cells and environments using singular value decomposition (SVD)—a method that decomposes high-dimensional datasets into orthogonal modes of variation ordered by statistical scale (**Methods**). In this representation, the first principal component (PC1) captures the largest scale of coordinated variation in the data, PC2 the second largest, and so on, producing a spectrum of PCs (‘eigenspectrum’) that describes how transcriptional variability is distributed across scales. We computed the eigenspectrum of the full expression matrix (cells × genes) and found that the top 30 PCs together captured 11.59% of the total transcriptional variance. Importantly, the variance explained by these PCs decreased approximately linearly with PC rank (**Fig. 1E**). Recent work has suggested that such scaling behavior indicates the presence of structured biological signals across multiple statistical scales rather than a small number of dominant programs [Zaydman et al., 2022a,b, Doran et al., 2025]. This observation suggested that meaningful transcriptional variation might extend well beyond the shallowest PCs.

To determine what biological information was encoded across these scales, we asked how environmental cues mapped onto different regions of the eigenspectrum. Each cell contributes to each PC to a certain degree, and each cell was part of a population exposed to a set of environmental stresses defining its complex environment (carbon source, osmolarity, temperature, media composition, and ROS stress). Standard dimensionality-reduction visualizations such as UMAP compress information across PCs and therefore obscure how information is distributed across statistical scales. To instead directly interrogate the eigen-spectrum, we applied a technique called SCALES (Spectral Correlation Analysis of Layered Evolutionary Signals) [Zaydman et al., 2022a,b, Doran et al., 2025]. SCALES quantifies how biological metadata maps onto different spectral layers of high-dimensional datasets in a purely data-driven manner. Briefly, SCALES computes the mutual information between transcriptional covariance patterns defined by windows of PCs and discrete environmental variables (**Methods**; **fig. S4**). The output of SCALES would therefore indicate where information about environmental cues lie across the spectrum of PCs.

Applying SCALES to sequential windows of PCs revealed two key features of transcriptional organization across the environments we studied. First, the shallowest PCs contained partial information about multiple environmental variables. Inspection of the genes with the highest loadings on these components showed that PC1 was enriched for glycolytic and gluconeogenic enzymes reflecting carbon source identity (e.g. *ENO2*, *PDC1*, *ADH1*), and PC2 for canonical stress-response genes involved in metabolism (e.g. *HSP12*, *HSP26*, *SSA1*) (Table S2). Thus, the leading modes collectively captured a broad, condition-agnostic transcriptional shift resembling the canonical ‘environmental stress response’ (ESR) [Gasch et al., 2000, Gasch and Werner-Washburne, 2002, Brauer et al., 2008]. Second, each environmental cue was resolved within distinct regions of the eigenspectrum spanning shallow to deeper statistical scales. Carbon source information appeared within the shallowest PCs, followed sequentially by media composition, osmolarity, temperature, and ROS stress (**Fig. 1F**). Importantly, shuffled controls in which environmental labels were permuted across cells produced mutual information values well below those of the true data across all PC windows, confirming that the observed hierarchy reflects genuine environmental signal rather than statistical noise (**Fig. 1F**, **fig. S4**). In other words, when viewed through the decoder of SVD, the ensemble of transcriptomes resolved into a spectrally ordered representation of environmental variables, with different cues occupying distinct statistical scales.

Together, these analyses revealed that transcriptional adaptation is distributed across separable spectral scales: a shared ESR-like program dominates the leading modes of variation, while distinct environmental cues emerge in progressively deeper components in an ordered manner. This structure suggested that environmental information is hierarchically organized within the yeast transcriptome. However, these analyses were performed on populations that had already reached steady-state growth in each environment. We therefore next asked whether this statistical hierarchy reflects a physiological hierarchy that emerges during the dynamics of adaptation.

### Statistical hierarchy of transcriptional variation is reflected in the dynamics of adaptation

We focused experiments on yeast populations adapting to changes in osmolarity and temperature—a combination of cues we did not assay within our initial dataset. Because osmolarity ranked above temperature in our statistical hierarchy, our model suggested that under combined stress, cells would mount the osmotic-stress program preferentially over the heat-shock program (HSR) (**Fig. 2A**) [Brewster and Gustin, 2014, Capaldi et al., 2008, Muzzey et al., 2009, Mettetal et al., 2008, Hohmann, 2002, Saito and Posas, 2012]. To test this hypothesis, we transcriptionally profiled cell populations via bulk RNA-seq while systematically varying the order and combination of osmotic and temperature challenges (**Fig. 2B**). We grew cells to exponential phase in rich media at 30°C and: (i) left them untreated; (ii) heat shocked them at 39°C; (iii) osmotically shocked them at 0.8 M NaCl; or (iv) shocked them with both 39°C and 0.8 M NaCl. Compared to heat shock only, genes associated with the heat shock response (HSR target genes) showed significantly lower induction in the dual stress condition (**Fig. 2C**, red data points) (**Methods**) [Zheng et al., 2016, Krakowiak et al., 2018, Pincus et al., 2018, Solis et al., 2016]. By contrast, the genes associated with osmotic shock (HOG target genes) were induced nearly to the same extent in hyperosmotic shock and in the dual stress condition (**Fig. 2C**, light blue data points) [Capaldi et al., 2008, Brewster and Gustin, 2014]. Moreover, the canonical HSR target gene, *HSP104*, showed the same induction in the dual stress as in osmotic shock, while the canonical HOG target gene, *STL1*, showed substantially increased induction in the dual stress compared to heat shock (**Fig. 2D**). Thus, osmotic stress prevented heat stress from inducing *HSP104*, but heat stress did not prevent osmotic stress from inducing *STL1*. Ribosomal protein genes (RPGs) were repressed under both heat and hyperosmotic stress, but their expression level in the dual stress more closely resembled hyperosmotic shock than heat shock (**Fig. 2E**) [Marion et al., 2004, Warner, 1999, Shore et al., 2021]. Across HSR targets, HOG targets, and RPGs, the measured expression levels in the dual stress condition deviated from the expected value of expression levels if heat shock and osmotic shock influenced gene expression independently (**Fig. 2C–E**, yellow data points). Considering the whole yeast transcriptome, the dual stress condition was better correlated with the osmotic stress condition (r=0.83) than with heat shock (r=0.45) (**fig. S5A**).

**Fig. 2:**
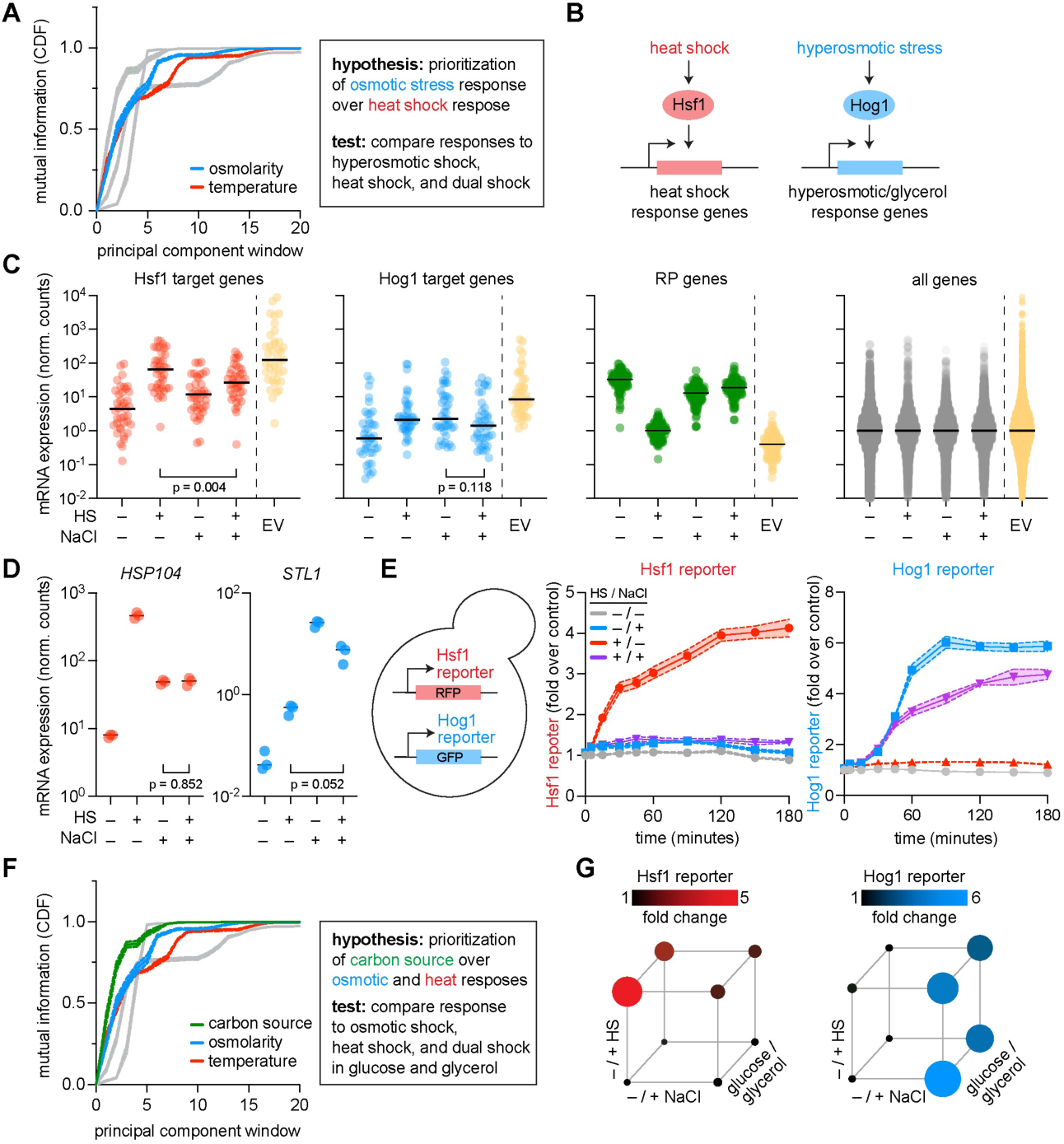
Hierarchical stress prioritization revealed within dynamics of adaptation to combined environmental perturbations. **A.** Mutual information (y-axis) across principal component windows predicts that osmotic stress is prioritized over heat shock (box). **B.** Hsf1 and Hog1 are master regulators of the heat shock response and osmotic stress response, respectively. **C.** Normalized mRNA expression of Hsf1 target genes, Hog1 target genes, ribosomal protein (RP) genes, and all genes across conditions with heat shock (39°C), osmotic shock (800 mM NaCl), or dual stress. The expected value (EV, dashed line) represents additive independent contributions of each stress. Horizontal bars indicate medians. **D.** Normalized mRNA expression of canonical Hsf1 target gene *HSP104* and Hog1 target gene *STL1* across the four conditions. **E.** Transcriptional fluorescent reporter measurements following heat shock (39°C), osmotic shock (0.8 M NaCl), or dual stress. Under dual stress, Hsf1 targets and Hsf1 reporter activity are suppressed relative to heat shock alone, whereas Hog1 targets and Hog1 reporter activity remain robustly induced. **F.** Extension of the hierarchical prioritization scheme to include carbon source. **G.** Transcriptional and fluorescent reporter measurements following heat shock (39°C), osmotic shock (0.8 M NaCl), or dual stress at 30 minutes when cells are grown in 2% glucose (front cube face) versus 2% ethanol/glycerol (back cube face, fig. S5). Non-stress condition is marked by the bottom left vertices. The area of each circle corresponds to the mean value of reporter fold change (*n*=3).

As an orthogonal method to measure induction of the HSR and HOG pathway, we measured the fluorescence of specific reporters (**Fig. 2E**, schematic) [Zheng et al., 2016, Stewart-Ornstein et al., 2012, Hao and O’shea, 2012]. The HSR reporter exhibited minimal induction in dual stress compared to heat shock (**Fig. 2E**, left graph). By contrast, the HOG reporter was robustly induced in the dual stress condition (**Fig. 2E**, right graph). When cells were given time to initiate the HSR prior to osmotic stress, subsequent addition of osmoshock still caused HSE-RFP fluorescence to plateau (**fig. S5B**). HSR inhibition was dose-dependent on both KCl concentration and temperature (**fig. S5B**). These results were consistent with our bulk transcriptional profiling data, with prioritization of the HOG pathway even more apparent in the reporter assays. Overall, these data demonstrate environmental epistasis between heat and osmotic shock at the level of gene expression, with the hyperosmotic shock program showing prioritization over the heat shock program—a result consistent with the statistical hierarchy we observed.

Next, we extended our validation of the statistical hierarchy an additional layer. The statistical hierarchy suggested that cells would prioritize their response to alternative carbon source over both the HSR and HOG pathways (**Fig. 2F**) [Visser et al., 1990, Merico et al., 2007]. Therefore, we used the reporter assays to test this by repeating the environmental epistatic cycle (−/+ heat shock, −/+ osmotic shock) with cells growing either in glucose or ethanol + glycerol (**Methods**). Cells grown in ethanol + glycerol showed reduced output of both the HSR and HOG pathways compared to cells grown in glucose (**Fig. 2G**, **fig. S5C**). Thus, the output of the HSR in response to heat shock depended both on carbon source and osmolarity, and the output of the HOG pathway depends on carbon source. These results illustrated a three-way environmentally-associated transcriptional epistasis consistent with the prioritization scheme suggested by the statistical hierarchy.

Overall, we found that the hierarchy derived from statistical analysis conducted across an ensemble of adapted yeast populations was reflected in how individual cell populations deploy stress responses when individual stressors are combined. Motivated by these results, we next asked whether this prioritization logic was encoded at the biological scale of single cells—as opposed to emerging at a population level—and if so, what molecular mechanisms allow one environmental cue to suppress adaptation to another cue in combined conditions.

### Adaptive prioritization within single cells occurs through global control of translation initia-tion

The prioritization of adaptation to osmoshock over heat-shock suggested the existence of a cross-pathway inhibition mechanism (**Fig. 3A**) [McClean et al., 2007, Madhani and Fink, 1997, Hao et al., 2013]. The master regulators of both pathways, Hog1 and Hsf1, accumulate in the nucleus upon activation [Brewster and Gustin, 2014, Hao and O’shea, 2012, Zheng et al., 2016, Sorger and Pelham, 1988]. Using yeast strains in which the only copies of Hog1 and Hsf1 are tagged in the genome with fluorescent proteins, we imaged cells across the conditions of heat shock, osmoshock, and dual stress (**Fig. 3B**) (**Methods**). Hog1 nuclear localization increased upon both osmotic shock and dual heat and osmotic shock, albeit to a lesser extent in the dual stress (**Fig. 3C**, light blue data points) [Muzzey et al., 2009, Vidal et al., 2013]. In contrast, Hsf1 nuclear localization increased upon heat shock but not in the dual stress condition (**Fig. 3C**, red data points) [Zheng et al., 2016]. Thus, osmotic stress prevented heat shock dependent nuclear enrichment of Hsf1, implicating upstream regulation of Hsf1 in the cross-pathway inhibition mechanism.

**Fig. 3:**
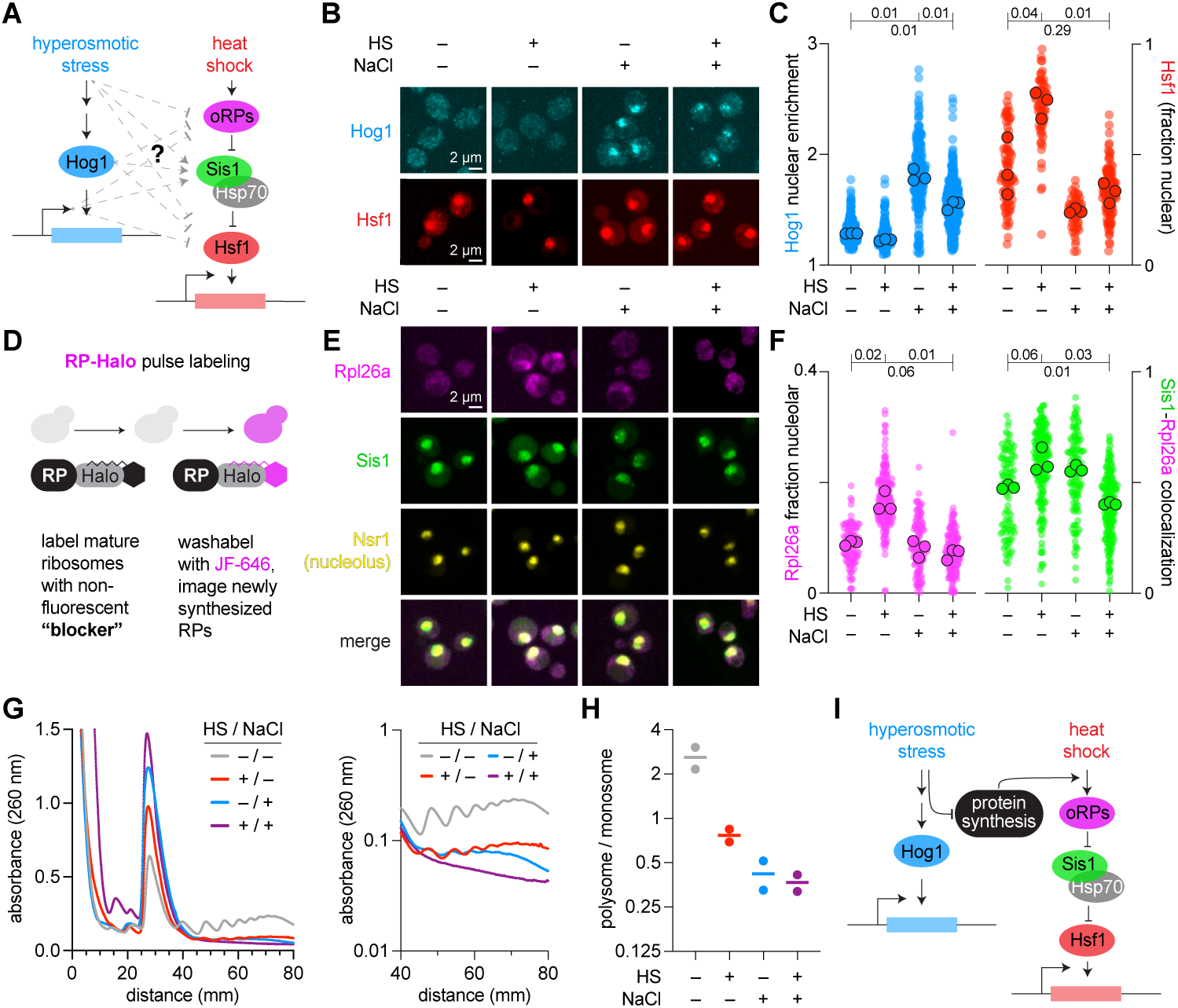
Mechanism of adaptive prioritization involves differential translation initiation. **A.** Schematic of cross-talk between osmoshock and heat shock pathways. **B.** Cellular localization of Hog1 and Hsf1 in the context of osmoshock (NaCl), heat shock (HS), or both stresses simultaneously. **C.** Nuclear enrichment of Hog1 and Hsf1 across the same set of environmental stresses in panel B. **D.** Schematic of HaloTag-based pulse labeling of Rpl26a. **E, F.** Imaging (E) and nuclear localization measurements of Rpl26a, Sis1, Nsr1, and Sis1–Rpl26a colocalization (F). **G.** Polysome profiles across stress conditions (left: linear scale; right: log scale). **H.** Polysome-to-monosome ratios across environmental conditions described in panel B. **I.** Schematic model for prioritization of osmoshock over heat shock response: osmotic stress inhibits orphan ribosomal protein translation.

Our results motivated a natural question: why is nuclear localization of Hsf1 dependent on the presence or absence of osmotic stress? In the absence of stress, it is known that the chaperone Hsp70 and co-chaperone Sis1 prevent Hsf1 from oligomerizing, binding DNA, and recruiting requisite transcriptional machinery for adaptation to heat shock [Krakowiak et al., 2018, Zheng et al., 2016, Garde et al., 2023, Ali et al., 2023, Garde et al., 2024, Shi et al., 1998, Zou et al., 1998, Kmiecik et al., 2020, Kmiecik and Mayer, 2022]. Heat shock and other stressors disrupt ribosome biogenesis and pre-rRNA processing, leading to the accumulation of unassembled “orphan” ribosomal proteins (oRPs) [Ali et al., 2023, Tye et al., 2019, Albert et al., 2019]. oRPs condense and recruit Sis1 and Hsp70 to the nucleolus and away from Hsf1, leaving Hsf1 free to activate the HSR genes [Ali et al., 2023, Feder et al., 2021, Frottin et al., 2019]. We therefore hypothesized that osmotic shock may interact with the capacity for oRPs to recruit Sis1 and Hsp70 away from Hsf1. To test this idea, we used HaloTag-based pulse labeling of Rpl26a, a ribosomal protein of the large subunit, to monitor oRP accumulation (**Fig. 3D**) (**Methods**) [Ali et al., 2023]. We imaged newly synthesized Rpl26a in cells also expressing fluorescently tagged Sis1 and the nucleolar marker Nsr1 across the heat shock, osmoshock, and dual stress conditions (**Fig. 3E**). Pulse-labeled Rpl26a showed increased nucleolar localization upon heat shock but not in dual heat and osmotic shock (**Fig. 3F**, magenta data points). Likewise, the increase in co-localization of Sis1 with pulse-labeled Rpl26a upon heat shock was abrogated in the dual stress condition (**Fig. 3F**, green data points) [Feder et al., 2021, Klaips et al., 2020]. These data demonstrated that osmotic shock prevented accumulation of oRPs and co-localization of Sis1 upon heat shock, explaining why HSR induction is reduced in the dual stress compared to heat shock.

We next interrogated why osmoshock prevented accumulation of oRPs while heat shock did not. Production of ribosomal proteins requires ongoing protein synthesis, and full activation of the HSR requires that cells be actively translating proteins prior to heat shock [Tye and Churchman, 2021, Ali et al., 2023, Masser et al., 2019]. We assessed global protein synthesis by monitoring polyribosome (polysome) abundance—clusters of ribosomes simultaneously translating a single mRNA, whose levels reflect the rate of translation initiation— across the cycle of heat and osmoshock (**Methods**) [Aboulhouda et al., 2017]. Compared to unstressed cells, all three stress conditions resulted in reduced polysome and increased monosome peaks, indicating that these stresses result in an inhibition of translation initiation (**Fig. 3G**, left panel) [Costa-Mattioli and Walter, 2020, Iserman et al., 2020, Cherkasov et al., 2013, Sonenberg and Hinnebusch, 2009]. However, although doubly shocked cells showed a complete collapse of polysomes, osmotically shocked cells retained low levels of light polysomes, and heat shocked cells retained low levels of both light and heavy polysomes (**Fig. 3G**, right panel). The ratios of polysome to monosome peaks show that the dual stress condition more closely resembles osmotic shock than heat shock (**Fig. 3H**). Together, these results demonstrate the mechanism for crosstalk inhibition between the HOG pathway and the HSR: osmotic shock results in the collapse of polysomes, preventing the accumulation of oRPs and other newly synthesized proteins, thereby leaving Sis1 and Hsp70 available to continue to repress Hsf1 from activating the HSR (**Fig. 3I**) [Ali et al., 2023, Krakowiak et al., 2018, Garde et al., 2024].

Taken together, our findings demonstrate that prioritization is not merely a population-level averaging phenomenon but a property molecularly encoded within single cells. Under dual stress, individual cells show robust Hog1 activation but diminished Hsf1 nuclear enrichment. The associated reduction in oRP accumulation and stronger collapse of polysomes in dual stress illustrates differential translation initiation as a control point that can gate pathway output [Costa-Mattioli and Walter, 2020, Pakos-Zebrucka et al., 2016, Iserman et al., 2020]. Although we focused on a particular set of stress programs, our findings offer mechanistic resolution for how the statistical hierarchy of adaptive prioritization is directly instantiated at the level of single cells.

### Prolonged evolution under constant stress reorganizes adaptive prioritization

As our results demonstrated that adaptive prioritization was encoded within the molecular biology of single cells, we next asked to what degree is the hierarchical architecture of prioritization an intrinsic property of cellular organization or contingent on the history of selection and variation that a species faces? As it is not possible to go back in time and profile yeast populations, we addressed this problem using forward evolution via continuous culture [Dhar et al., 2013, Caspeta and Nielsen, 2015, Dunham et al., 2002, Lang et al., 2013, Levy et al., 2015]. Our hypothesis was that if the hierarchical structure of adaptability was generated through a ‘hard-wired’ design principle, it would be preserved even under prolonged selection for a single stress. Conversely, if its origin depended on a history of exposure to environmental heterogeneity, sustained evolution in a constant condition may reorganize the hierarchical prioritization of adaptation.

We used a laboratory evolution setup, the eVOLVER system [Wong et al., 2018], to maintain yeast cell populations in continuous culture in logarithmic growth phase for more than nine months (> 3000 generations) and selected for long-term fitness in synthetic media with 0.8 M KCl (**Fig. 4A; fig. S6A**) [Basan et al., 2020, Erickson et al., 2017, Scott et al., 2010, Klumpp et al., 2009]. This process yielded a strain we term the ‘derived strain’. Here we focus on a single deeply characterized derived lineage to establish the principle that prolonged selection reorganizes adaptive prioritization. Replicate evolution experiments across multiple selective environments, which confirm and extend these findings, will be reported in a companion study (Lan et al.). The derived strain showed increased maximal growth rate relative to the ancestral strain in synthetic media in 0.8 M KCl and retained its ability to grow as well as the ancestral strain in synthetic media lacking KCl (**fig. S6B**). However, the derived strain grew more slowly than the ancestral strain in the other 18 conditions, with a loss of diauxic growth in glucose conditions, and a loss of viability at elevated temperature with ethanol + glycerol as a carbon source (**fig. S6B**). Competitive growth assays showed that the derived strain had reduced fitness compared to the ancestral strain in all conditions tested except synthetic media with 0.8 M KCl, the condition in which it was selected (**Fig. 4B**).

**Fig. 4:**
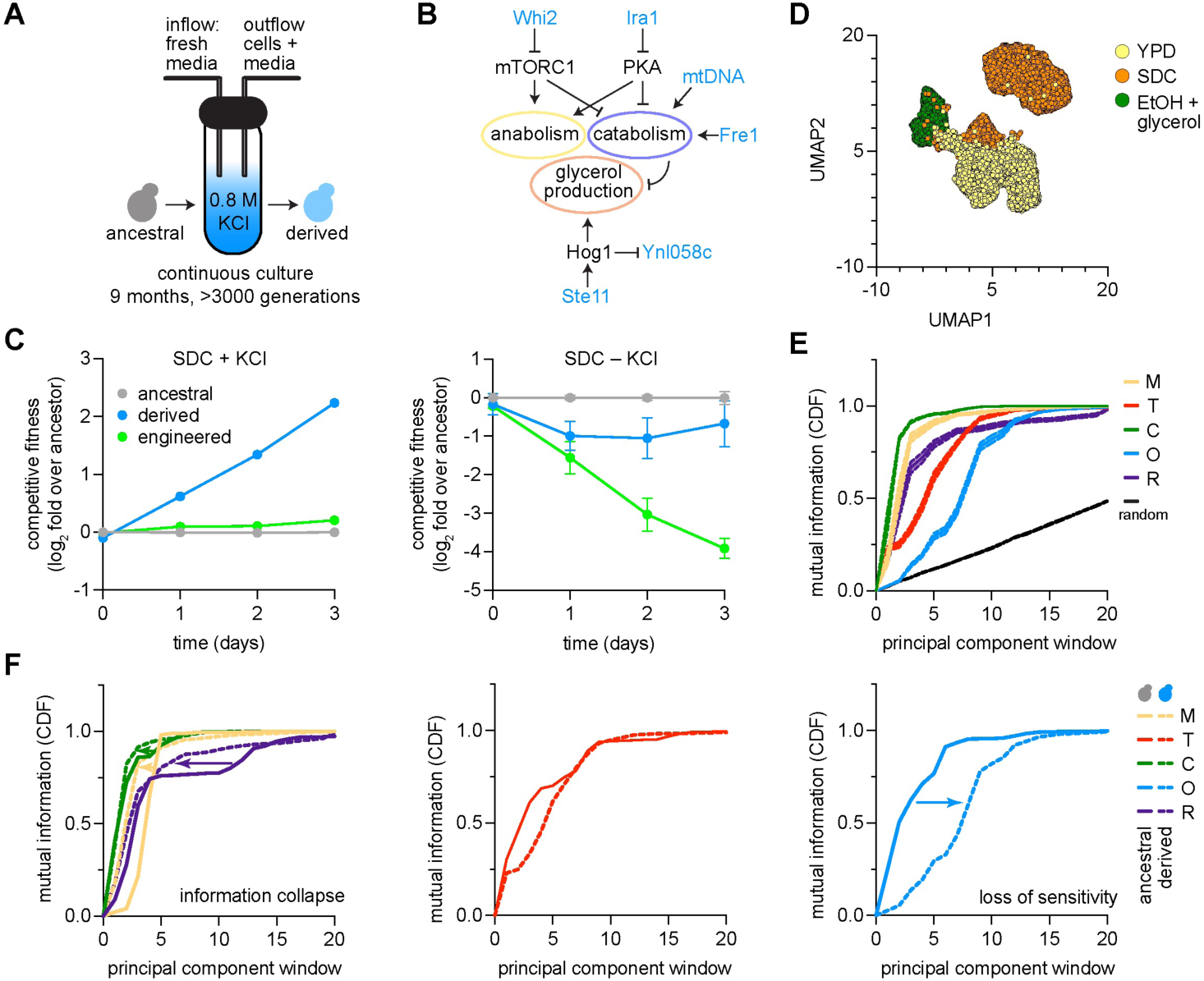
Long-term evolution under constant osmotic stress reorganizes adaptive prioritization. **A.** Schematic of laboratory evolution experiment. Ancestral yeast populations were maintained in continuous culture in synthetic media with 0.8 M KCl for >3,000 generations (∼9 months; eVOLVER system) yielding a ‘derived’ strain. **B.** Model associating mutations observed in derived strain with effect on osmotic stress response. **C.** Competitive fitness assays between ancestral, derived, and ‘engineered’ strain—a strain engineered with mutations in genes shown in panel B. **D.** UMAP of single-cell transcriptional profiles for derived cells (dots). Derived cells were exposed to all environments in Fig. 1A; cells are colored by specific environmental cues (YPD, SDC, and ethanol plus glycerol). **E.** Mutual information (MI; cumulative density) between environmental cues (color key) and eigenspectrum of transcriptional covariation for all cells shown in panel D (x-axis). ‘Random’ reflects null expectation of MI. **F.** Comparison of MI curves for ancestral and derived strains.

Genome sequencing of the derived strain revealed the presence of termination codons in protein coding sequences of five genes (Table S3): *FRE1*, encoding a high-affinity iron transporter; *IRA1*, encoding a negative regulator of Ras signaling; *STE11*, encoding a positive regulator of the HOG pathway; *WHI2*, encoding a cell cycle checkpoint protein and negative regulator of mTORC1 signaling; and *YNL058C*, encoding a membrane protein repressed by HOG signaling (**Fig. 4C, fig. S7A, B**) [Chen and Thorner, 2007, Brewster and Gustin, 2014, Turco et al., 2023]. In addition to these genomic mutations, the derived strain harbored reduced levels of mitochondrial DNA compared to the ancestral strain and unchanged ribosomal DNA content (**fig. S7C, D**). Given the known roles of Ras and mTORC1 in regulating growth [Prouteau et al., 2017, Delarue et al., 2018], and the HOG pathway in arresting the cell cycle during osmotic stress [Brewster and Gustin, 2014, Capaldi et al., 2008], we hypothesized these mutations may be responsible for the increased proliferation of the derived strain in hyperosmotic conditions. To evaluate their sufficiency for generating the phenotypes observed in the derived strain, we engineered these mutations into the ancestral strain by serial genome editing with CRISPR/Cas9 (**Methods**) [Bao et al., 2015]. However, rather than recapitulating the increased fitness of the derived strain relative to the ancestral strain in 0.8 M KCl, competitive fitness assays revealed no difference in fitness in the CRISPR/Cas9 engineered strain relative to the ancestral strain in 0.8 M KCl and substantially reduced fitness relative to the ancestral strain in the absence of KCl (**Fig. 4B**). These mutations were therefore insufficient to improve fitness in KCl, suggesting that mitochondrial DNA differences, epigenetic changes, specific mutation acquisition orders, or other forms of epistasis were also required to specify the phenotype of the derived strain.

We therefore asked whether prolonged selection under constant osmotic stress had altered the broader architecture of adaptation. In the derived strain, we repeated profiling by scRNA-seq in 18 of the 20 environments we assayed for the ancestral strain (the derived strain failed to grow in SC or YP media with ethanol + glycerol at elevated temperature as mentioned). We grew cultures of the derived strain to maximal logarithmic phase growth rate in each environment, harvested cells for scRNA-seq, and sequenced ≥ 1000 cells with ≥ 500 genes expressed from each sample (**fig. S8**, Table S1) [Luecken and Theis, 2019, Sarkar and Stephens, 2021]. UMAP revealed separation of cells grown in ethanol + glycerol compared to cells grown in glucose and separation within the glucose cluster of cells grown in YP versus SC media (**Fig. 4D**, **fig. S9**). We then computed the mutual information between PCs encoding patterns of transcriptional variation across cells within all 18 environments and environmental variables [Zaydman et al., 2022a]. Relative to the ancestral strain, the derived strain exhibited a differential redistribution of information across the eigenspectrum of transcriptional variation (**Fig. 4F**). Information corresponding to media composition, carbon source, and reactive oxygen species collapsed toward shallower spectral windows, consistent with compression of environmental information into dominant modes of variation (**Fig. 4E**, left). Temperature sensitivity remained largely unchanged with information distributed across a similar range or spectral windows as in the ancestor (**Fig. 4E**, middle). Information associated with osmotic stress shifted toward deeper spectral windows, indicating a loss of prominence relative to the dominant modes of transcriptional variation (**Fig. 4E**, right).

Because the statistical structure of environmental information in the derived strain suggested that multiple cues are encoded within a shared set of dominant modes, we next sought to interrogate what biological program occupies these modes. In principle, given selection in high osmotic stress, the shallowest modes of transcriptional variation could incorporate osmotic stress into a generalized response or exclude it, reflecting decoupling from the shared generic stress axis. To distinguish these possibilities, we examined the gene loadings associated with the leading eigenmodes of transcriptional variation. Strikingly, the genes with the highest loadings on these dominant modes in the derived strain remained canonical ESR genes, indistinguishable in identity from those in the ancestral strain. PC1 was dominated by canonical repressed ESR genes that are primarily glycolytic enzymes—including *ENO2*, *FBA1*, *CDC19*, *GPM1*, *TPI1*, and *PDC1* (Table S4). Moreover, loading magnitudes were relatively uniform across these genes, indicating that no single functional subprogram dominated the leading mode. Thus, rather than incorporating osmotic stress into a generalized program, the derived strain exhibited convergence of multiple environmental responses—particularly carbon, media, and ROS—onto a shared generic stress axis corresponding to the shallowest layer of the ancestral hierarchy. In contrast, osmotic stress was not represented within these dominant modes, consistent with its decoupling from the generic stress program. As a result, while derived cells retained the ability to detect stress, their capacity to resolve and integrate specific environmental cues was reduced.

Together, these results show that the hierarchical architecture of adaptive prioritization is neither fully hard-wired nor uniformly eroded under sustained selection but is instead selectively reorganized by evolutionary history. Prolonged evolution in a constant environment redistributes environmental cues within this hierarchy—compressing some into shared dominant modes while selectively deprioritizing the very environmental variable under sustained selection, here osmotic stress, as an organizing axis of the response. These findings suggest that sustained selection does not amplify the representation of an environmental variable, but instead converts it from a regulated input into a constitutive condition of the system.

### Loss of stress integration in the derived strain

The statistical structure of transcriptional variation in the derived strain illustrated that long-term evolution selectively deprioritized osmotic stress while preserving temperature as a distinct axis of transcriptional variation. This raised the possibility that the derived strain would retain responses to individual stresses, but exhibit defects in integrating osmotic stress with other environmental cues. Because transcriptional response to temperature remained spectrally distinct while osmotic stress was selectively deprioritized in the derived strain, the combination of these stresses provided a direct test of whether the derived strain could integrate a distinct environmental stress (heat shock) with one that was placed into the background by prolonged selection (osmoshock). We therefore compared transcriptional responses, translational dynamics, and growth of ancestral and derived strains under single and combined heat and osmotic stress conditions.

We grew the derived cells to exponential phase in rich media at 30°C and: (i) left them untreated; (ii) heat shocked them at 39°C; (iii) osmotically shocked them at 0.8 M NaCl; or (iv) shocked them with both 39°C and 0.8 M NaCl. Under the unstressed condition or following treatment with 0.8 M NaCl, the transcriptome of the derived cells was largely unchanged compared to the ancestral strain (*R*^2^=0.97 in both conditions), with both HSR and HOG genes expressed equally in the two strains (**Fig. 5A**, top left and lower left). Notable differences in the basal transcriptome of the derived strain compared to the ancestral strain include reduced expression of pheromone response genes, consistent with its mutation in *STE11* [Chen and Thorner, 2007, Madhani and Fink, 1997], and increased iron transport, consistent with its *FRE1* mutation (**fig. S7**). Following heat shock, the derived strain robustly induced the HSR but did not repress ribosomal protein gene expression to the same extent as the ancestral strain, resulting in a reduced transcriptome-wide correlation (*R*^2^=0.92) (**Fig. 5A**, top right) [Marion et al., 2004, Shore et al., 2021]. In the dual stress condition, the derived strain failed to fully induce either the HSR or the HOG target genes, resulting in a reduced correlation with the ancestral strain (*R*^2^=0.83) (**Fig. 5A**, bottom right). Whereas expression of the canonical HSR and HOG target genes *HSP104* and *STL1* exemplified the prioritization of the HOG pathway over the HSR in the ancestral strain, neither gene was induced in the derived strain to the same extent in the dual stress as in its cognate single stress nor to its level in the dual stress in the ancestral strain (**Fig. 5B**). Thus, rather than prioritizing one response over the other, the derived strain induced neither the HSR nor the HOG pathway in the dual stress condition (**fig. S10**).

**Fig. 5:**
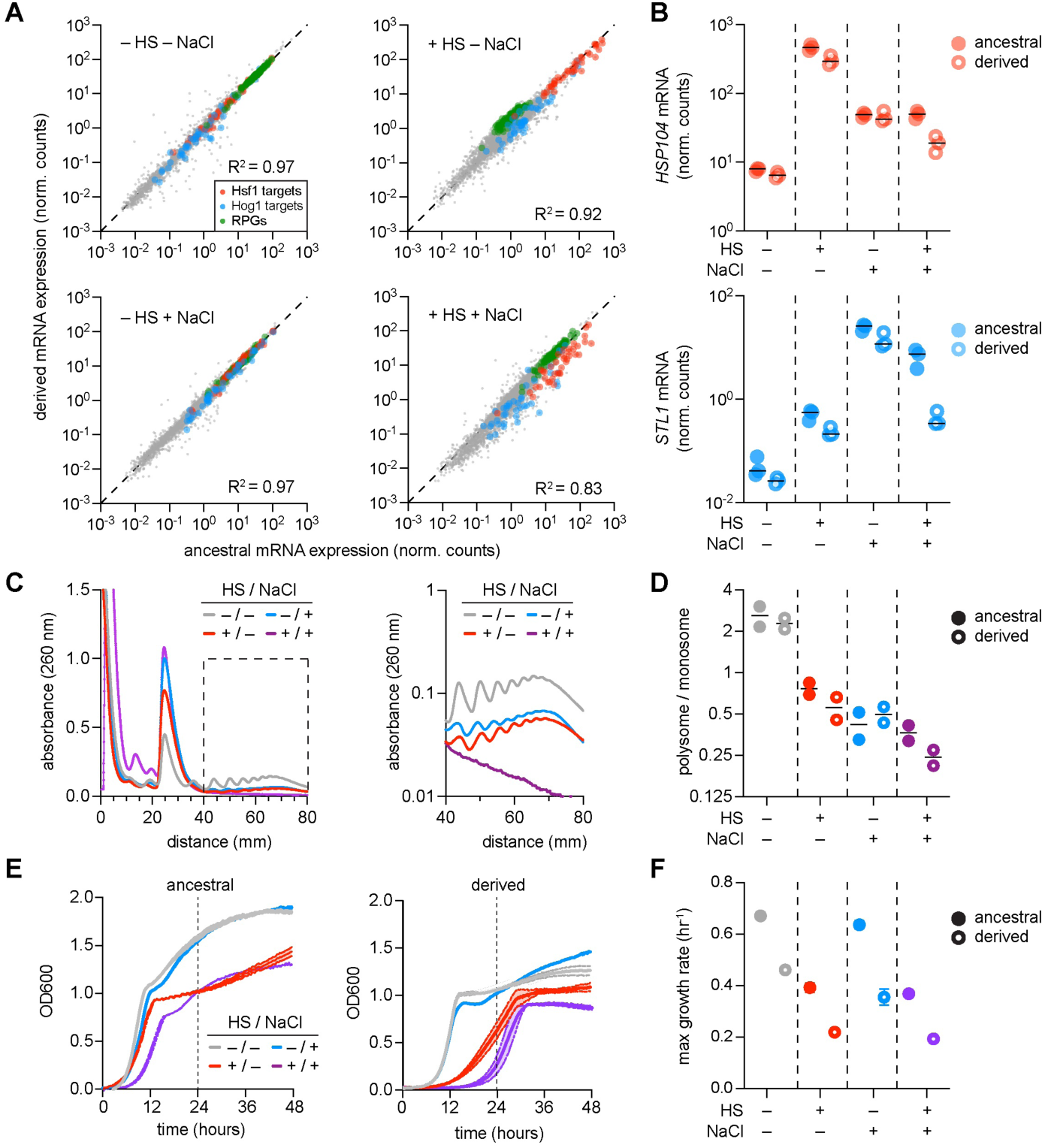
Examining the loss of prioritization between osmotic shock and heat shock in the derived strain. **A.** Scatter plots comparing transcriptome-wide mRNA expression (normalized counts) between ancestral and derived strains across four conditions: −HS/−NaCl, +HS/−NaCl, −HS/+NaCl, and +HS/+NaCl. Hsf1 targets (red), Hog1 targets (blue), and ribosomal protein genes (RPGs, gray) are highlighted (*R*^2^ values indicated for each condition). **B.** Expression of *HSP104* (top) and *STL1* (bottom) across the four stress conditions in ancestral (filled circles) and derived (open circles) strains. **C.** Polysome profiles of the derived strain across heat shock and osmotic shock conditions (left: linear scale; right: log scale). Compare to ancestral profiles in Fig. 3G. **D.** Polysome-to-monosome ratios for ancestral (filled) and derived (open) strains across the four stress conditions. **E.** Growth curves (OD_600_) of ancestral (left) and derived (right) strains across the four conditions over 48 hours. **F.** Maximum growth rates for ancestral (filled) and derived (open) strains across the four stress conditions.

The inability to induce either pathway in the dual stress conditions suggested that the derived cells were unable to adapt to the dual stress. Indeed, at the level of global protein synthesis, the derived strain showed substantially lower ratios of polysomes to monosomes than the ancestral strain in the dual stress condition—despite showing an increase in polysome content relative to the ancestral strain in osmotic stress alone (**Fig. 5C, D**) [Cherkasov et al., 2013]. At the level of growth, the loss of adaptability in the derived strain was evident across osmoshock, heat shock, and dual stress. Although the derived cells exhibited a higher maximum growth rate compared to the ancestral strain in osmotic stress, it showed (i) an inability for diauxic growth under nonstress and osmotic stress conditions, (ii) reduced maximum growth rate at elevated temperature, and (iii) the inability to effectively grow in the dual stress condition (**Fig. 5E, F**). These data demonstrate that, in evolving to grow in osmotic stress conditions, the derived strain lost the capacity to integrate dual stresses into a coherent adaptive response thereby leading to a drastic loss in fitness.

### Evolutionary history is a primary regulator of adaptability

The reconfiguration of the environmental hierarchy in the derived strain suggested that long-term selection under constant stress rewired transcriptional regulation in a global manner. To contextualize the transcriptional rewiring due to long-term evolution relative to that of adapting to different environmental conditions, we merged the scRNA-seq datasets collected for the ancestral and derived cell populations across all environmental conditions. Analysis of this merged, single dataset would characterize the relative importance of evolutionary history as its own ‘environmental cue’ as compared to the other environmental cues within our experimental workflow.

In a UMAP of the combined dataset, cells aggregated primarily by strain identity. Ancestral and derived strains formed distinct clusters whereas environmental separation was relatively diminished with the exception of carbon source (**Fig. 6A**). To evaluate this property in a more quantitatively rigorous manner, we computed the mutual information between the pattern of transcriptional variation across PCs with strain identity plus all other environmental cues (carbon source, osmotic environment, temperature, media composition, and presence/absence of reactive oxygen species). We found that the PCs harboring the highest data-variance—the leading eigenmodes—were enriched for information regarding environmental carbon source and strain identity (e.g. whether a cell was from the ‘ancestral’ versus ‘derived’ populations). In contrast, information about other environmental variables—osmotic environment, temperature, pH, and presence/absence of ROS—was displaced away from the shallowest eigenmodes and distributed across intermediate regions of the spectrum (**Fig. 6B**).

**Fig. 6:**
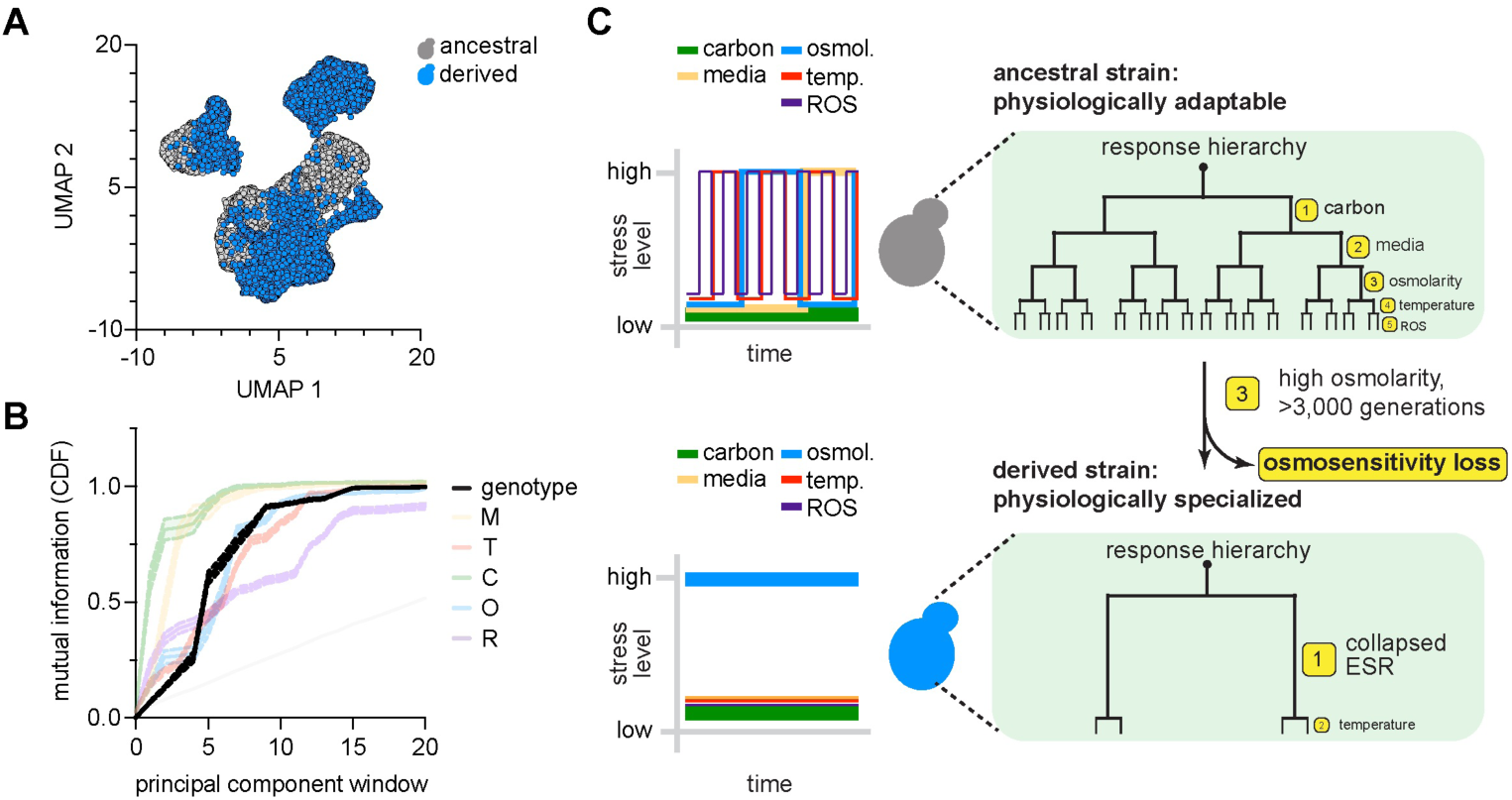
Ancestry is a primary regulator of adaptability. **A.** UMAP embedding of all cells profiled by scRNA-seq across all environments including ancestral and evolved strains. Each cell (dot) colored by whether the cell is ancestral (gray) or evolved (blue). **B.** Mutual information between various environmental cues, including ancestry, and structure of transcriptional covariation. **C.** Model of how evolutionary history affects the architecture of adaptability. Dynamically fluctuating selection history yields adaptive prioritization (top). Prolonged constant selection history deprioritizes adaptive response to selective pressure and emphasizes generalized stress response.

As the ancestral strain likely evolved under dynamically complex environmental cues, our results suggest a model where adaptability is an evolved property reflecting the statistical structure of environmental history. When organisms experience diverse and fluctuating conditions of selection, regulatory architectures are organized to preserve sensitivity across multiple environmental axes. In our case, this manifest in a hierar-chically distributed encoding of environmental information in the yeast transcriptional response (**Fig. 6C**, top). In contrast, prolonged exposure to a single dominant selective pressure compresses this hierarchy and deprioritizes response to the single pressure (**Fig. 6C**, bottom). In this way, the evolutionary history itself becomes a dominant axis of variation for the system, effectively acting as an additional ‘environmental variable’ that can supercede external conditions localized to physiological timescales.

## Discussion

Understanding how cells adapt to complex environments is a fundamental biological problem. Here, we found that (i) adaptability is hierarchically organized, (ii) this hierarchy emerges from regulatory dynamics within individual cells, and (iii) the architecture of adaptability is contingent on the history of evolutionary selection.

Classically, adaptation to stress has been conceptualized as a two-step nested model: cells first mount a conserved global stress response, then deploy environment-specific transcriptional programs [Gasch et al., 2000, Gasch and Werner-Washburne, 2002, Berry and Gasch, 2008]. In this view, adaptation assumes a ‘star-like’ geometry—each environment eliciting an idiosyncratic, largely independent branch from a common core. Our results support a different model. Rather than responding independently to each perturbation, cells decode complex environments into a hierarchy of response channels. Transcriptional adaptation is not purely bespoke—it is gated by a prioritization scheme in which some environmental dimensions dominate others. Adaptability therefore reflects hierarchically structured constraints rather than a collection of independent stress programs [Young et al., 2013, Stern et al., 2007]. Importantly, this hierarchy is not fixed. Sustained selection reorganizes the prioritization scheme itself [Dhar et al., 2013, Caspeta and Nielsen, 2015, Kvitek and Sherlock, 2013, Venkataram et al., 2016]. Thus, the more fundamental generative process governing the architecture of adaptability is not the molecular logic of any single stress-signaling pathway enabling current fitness, but rather the historical pattern of selection and variation. The evolutionary process sculpts the internal ordering of environmental demands. In this view, adaptability is an inherited molecular landscape that evolutionary pressures continually rewrite [Rutherford and Lindquist, 1998, Rohner et al., 2013, Cowen and Lindquist, 2005, Jarosz and Lindquist, 2010].

The collapse of the response hierarchy for various environmental cues in the derived strain speaks to a broader principle: that biological systems under perturbation often respond along surprisingly few axes of variation. In the ancestral strain, the leading eigenmode captures the canonical environmental stress response—genes such as *HSP26*, *SSA1*, *HSP12*, and *SOD2* that are induced broadly across conditions [Gasch et al., 2000]—while deeper eigenmodes separately encode information about carbon source, osmolarity, temperature, and other environmental features. The derived strain retains the same leading mode, with indistinguishable top-loading genes and markedly uniform loading magnitudes across canonical ESR and metabolic genes. What it has lost are the additional, environment-specific response programs that enable the ancestral strain to tell stresses apart outside of heat shock. This kind of low-dimensional compression—where diverse inputs are funneled into a single dominant response axis—has been formalized as the emergence of “soft modes” in biological systems: low-dimensional axes of variation that arise when regulatory networks are selected to maintain homeostasis against high-dimensional perturbations [Russo et al., 2025a,b]. Remarkably, an analogous phenomenon has been observed at the scale of organismal morphology where genetic and environmental perturbations to *Drosophila* wing development produce variation along a single dominant phenotypic mode rather than across many modes that the complexity of the system could theoretically permit [Alba et al., 2021, Kryazhimskiy et al., 2014]. Our results extend this picture from a temporal snapshot to an evolutionary trajectory: prolonged selection under reduced environmental complexity does not merely preserve the generic stress axis but actively eliminates certain condition-specific programs layered on top of it, thereby canalizing transcriptional responses into a single undifferentiated response.

Overall, our findings recast adaptability as an emergent, history-dependent architecture. By revealing the hierarchical structure of adaptability and that this structure can be reorganized by selection, our work opens a path towards a theory of flexibility within living systems [Freddolino et al., 2018, Stern et al., 2007, Guan et al., 2012]. An exciting future direction is to directly engineer the statistical history of environmental pressures with the goal of encoding and tuning adaptability into evolved systems. It is therefore possible that the findings described here could lay the foundation for engineering synthetic living systems with desired adaptive attributes.

## Materials and Methods

### Yeast strains and culture conditions

*Saccharomyces cerevisiae* strains used in this study are listed in Table S5. All strains are derived from the w303 parent strain.

Unless otherwise indicated, cells were grown at 30°C with shaking at 200 rpm in yeast extract–peptone–dextrose (YPD) medium. Overnight cultures were grown to saturation, diluted to an initial optical density (OD_600_) of 0.05 in the appropriate media, and allowed to grow to mid-logarithmic phase prior to experimentation. For stress or perturbation experiments, cells were transferred into condition-specific media for dilution as described below.

### Selection of environmental stressors and degree

We selected stress conditions that perturb distinct and well-characterized cellular programs in budding yeast while remaining within sublethal, physiologically interpretable ranges (Table S6). Comparison of rich medium (YP) and defined synthetic complete (SC) medium probes the degree of nutrient buffering and biosynthetic demand, distinguishing growth supported by abundant amino acids and peptides from growth requiring *de novo* synthesis. Temperature shift from 30°C (optimal growth) to 37°C induces a defined heat stress response and challenges proteostasis, membrane fluidity, and ribosome biogenesis. Carbon source substitution from fermentable D-glucose to non-fermentable glycerol/ethanol (2%/2%) enforces respiratory metabolism and mitochondrial dependence, thereby altering energy generation, redox balance, and transcriptional state. Osmotic stress was induced using 500 mM sorbitol (non-ionic osmolyte) and 500 mM KCl (ionic osmolyte) to distinguish general osmotic pressure effects from ion-specific stress signaling, particularly activation of the HOG pathway. Finally, oxidative stress was imposed using diamide (50 µM), which preferentially oxidizes thiol groups and perturbs redox homeostasis, and hydrogen peroxide (50 µM), which generates reactive oxygen species and activates canonical oxidative stress pathways. Together, these perturbations span nutrient, proteotoxic, metabolic, osmotic, ionic, and redox axes, enabling comparison of cellular responses across qualitatively distinct degrees and types of environmental stress.

### Stress time course via fluorescent reporter assays

Fluorescent transcriptional reporters were integrated at the *LEU* locus and expressed under the control of HSE or pHor2 to report on heat shock activity or osmotic stress activity, respectively.

Three biological replicates of each strain were serially diluted 5 times for a final dilution of 1:125 and grown in synthetic dextrose media overnight at room temperature. When cells had reached mid-log phase, 520 µL of each replicate were transferred to microcentrifuge tubes and aerated by shaking in a thermomixer at 30°C for one hour. Then, cells were either heat shocked, osmotic shocked, or doubly shocked. Cells were heat shocked by transferring the microcentrifuge tube into a preheated thermomixer set to 39°C. Cells were osmotic shocked by adding 125 µL of 4 M NaCl SDC, to a final concentration of 800 mM NaCl. At each time point, 50 µL of cells were transferred to a 96 well plate of SDC at 50 µg/mL final concentration cycloheximide. After time course collection, the plate was incubated at 30°C for 1 hour to promote fluorescent reporter maturation before cytometry.

### Flow cytometry

HSE-mApple and pHor2-GFP reporter levels in heat shock, osmotic shock, and double shock time course were measured with a BD Fortessa HTS 4-15 benchtop cytometer at the University of Chicago Cytometry and Antibody Technology Facility. Fluorescent measurements of 10,000 cells per replicate were performed using the 488-525 FITC fluorescence filter (pHor2-GFP) and 561-PE Dazzle (HSE-mApple). Raw fluorescence values were normalized by side scatter in FlowJo, and median fluorescence value was calculated for each of the three replicates.

### Quantitative growth assay

Parent or derived yeast strains were grown overnight at 30°C in YPD. In the morning, cells were diluted to OD_600_ of 0.05 in one of the 20 selected environmental configurations. 600 µL of culture was loaded into 48-cell plates and growth dynamics were measured using a Spectrostar Nano plate reader. OD_600_ measurements were recorded every 20 minutes for 48 hours with continuous shaking. Maximum growth rates were estimated by fitting a linear model to the log-transformed OD_600_ values during exponential growth.

### Fixed-cell imaging

Constructs containing fluorescently tagged Hog1, Hsf1, Sis1 and Rpl26a were used for imaging experiments (see Table S3 for combination of fluorophore tags). Cells were grown at 30°C with 200 rpm shaking overnight. In the morning, cells were diluted to OD_600_ of 0.1 and grown in the same conditions for 4 hours to reach log phase. Then, cells were either heat shocked, osmotic shocked, or doubly shocked. Cells were heat shocked by transferring 200 µL of cells into a preheated microcentrifuge in a thermomixer set to 39°C. Cells were osmotic shocked by adding 100 µL of 4 M NaCl SDC to 400 µL of cells to a final concentration of 800 mM NaCl. Reactions were terminated by addition of paraformaldehyde (16% stock) to a final concentration of 1% and fixed for 5 minutes at room temperature. Fixation was quenched by addition of glycine (1.25 M stock) to a final concentration of 100 mM. Cells were immediately plated onto Concanavalin A coated imaging 96-well plates for immobilization. Cells were then washed and imaged in KPIS media (1.2 M sorbitol, 0.1 M potassium phosphate, pH 7.5). Imaging was performed using a SoRa Marianas spinning disk equipped with a 100×/NA 1.3 oil objective and Yokogawa camera. Image processing and single-cell quantification were performed using CellQuant [Neferkara et al., 2026], an open-source Python pipeline for automated single-cell image analysis.

### Newly synthesized ribosome imaging

For HaloTag labeling experiments, 100 µL of log-phase culture was transferred to an Eppendorf tube and incubated in a thermomixer at 30°C with 1200 rpm shaking for 1 hour. For Rpl26a pulse–chase experiments, pre-existing ribosomes were blocked by adding 1 µL of 10 mM HaloTag blocker (100× stock) to 100 µL of culture (final 100 µM) and incubated for 15 minutes. Cells were pelleted by centrifugation for 30 seconds at room temperature, washed twice with 100 µL SDC to remove excess blocker, and resuspended in 20 µL SDC.

For stress and Halo dye labeling, 20 µL of cells were transferred into pre-prepared tubes containing 5 µL of 5 µM JF646 HaloTag ligand in SDC with the indicated stress conditions. Stress conditions included: no stress (tube at 30°C), osmotic stress (4 M NaCl SDC in tube), heat stress (tube pre-warmed to 39°C), and combined heat and osmotic stress (4 M NaCl SDC pre-warmed to 39°C). For heat-containing conditions, tubes were equilibrated at 39°C for at least 30 minutes prior to cell addition to ensure uniform temperature. Heat shock response (HSR) dynamics were assayed on the 2–10 minute timescale, whereas Hog1 pathway dynamics were assayed at 15–30 minutes.

Cell preparation and imaging was performed as described above in *Fixed-cell imaging*.

### Single-cell RNA sequencing

Cells were grown overnight in YPD to saturation, diluted to an OD_600_ of 0.05 in condition-specific media, and harvested at mid-log phase. Mid-log phase was verified by growing the cells in the Spectrostar Nano plate reader and monitoring for previously measured log-phase OD and time. Cells were harvested via filtration and washed twice with PBS to remove remaining media before collection with a cell scraper. Cells were zymolyase treated [Jariani et al., 2020] prior to droplet generation.

Libraries were generated using the Chromium Single Cell 3’ Kit v3.1 according to the manufacturer’s protocol. Sequencing was performed on a NovaSeq 6000 at the University of Chicago Functional Genomics Facility (RRID: SCR_019196) [Luecken and Theis, 2019, Sarkar and Stephens, 2021].

### Bulk RNA sequencing

For bulk RNA-seq experiments, cells were grown overnight in YPD, diluted to an OD_600_ of 0.05 in condition-specific media, and harvested at mid-log phase (OD_600_ = 0.4–0.6, approximately 5 hours post-dilution) by centrifugation. Cell pellets were washed and resuspended in cryopuck buffer (30% RNase-free glycerol, 20 mM HEPES-KOH pH 7.5, 120 mM KCl, 2 mM EDTA, 1 U/µL SUPERase•In RNase inhibitor), then flash-frozen dropwise into liquid nitrogen to form frozen beads. Beads were cryomilled using a mixer mill at 30 Hz for 1.5 minutes, repeated 5 times with liquid nitrogen rechilling between cycles, to produce cryopowder.

RNA was extracted by adding 350 µL TRI Reagent directly to cryopowder, followed by incubation at room temperature for 5 minutes. RNA was purified using the Zymo Direct-zol RNA Miniprep Kit (R2050) according to the manufacturer’s protocol, including on-column DNase I digestion. RNA was eluted in 30–50 µL RNase-free water and stored at −80°C. Library preparation and sequencing were performed at the University of Chicago Functional Genomics Facility (RRID: SCR_019196). Libraries were sequenced on an Illumina NovaSeq 6000.

Raw reads were aligned to the *S. cerevisiae* reference genome (sacCer3) using Bowtie2. Aligned reads were coordinate-sorted and indexed using SAMtools. Read counts were assigned to genes using featureCounts using gene annotations from the sacCer3 GTF. Count matrices were generated using both gene ID and gene name identifiers. Alignment and assignment rates were computed per sample from SAMtools flagstat output and the featureCounts summary file, respectively.

### Bioinformatic processing for single-cell RNA sequencing

Sequencing reads were processed using Cell Ranger 7.1.0 (10x Genomics) with –expect-cells 2000, –localcores 8, and –localmem 64. Reads were aligned to the *S. cerevisiae* S288C reference genome assembly R64-1-1 (Ensembl; INSDC assembly GCA_000146045.2, gene annotation updated October 2018) with gene annotation imported from the Saccharomyces Genome Database (SGD). Gene-level UMI counts were obtained from the filtered feature-barcode matrices output by Cell Ranger (filtered_feature_bc_matrix.h5). All downstream analyses were performed in Python using Scanpy and in R.

Quality control and filtering were performed using Scanpy. Cells were retained if they expressed between 200 and 4,000 genes and had at least 300 total UMI counts. Genes expressed in fewer than 10 cells across the dataset were excluded. Gene expression counts were normalized to 10,000 counts per cell, log-transformed, and stored prior to downstream analysis.

### Information-theoretic analyses for single-cell RNA sequencing

To characterize the statistical structure of transcriptional adaptation across environments, we applied Singular Value Decomposition (SVD) to the log-normalized, library-size-normalized gene expression matrix (unscaled, all-gene counts from adata.raw.X). Prior to SVD, highly variable genes were identified and expression values were scaled to unit variance across cells; SVD was then applied to the full log-normalized unscaled matrix. This procedure yields a set of left singular vectors (LSVs) ordered by the amount of transcriptional variance they capture, analogous to principal components.

To quantify which environmental variables were encoded within specific regions of the PC spectrum, we used the Spectral Correlation Analysis of Latent Environmental Signals (SCALES) method, developed in Zaydman et al. [2022a] and implemented in the spipy package (aramanlab/spipy). Briefly, LSVs were divided into sliding windows of 3 consecutive components beginning at PC1. For each window, cell–cell spectral correlations were computed as the inner product of each cell’s projection onto that window of LSVs. Mutual information (MI) was then calculated between the distribution of spectral correlations and a binary label indicating whether each pair of cells shared the same value for a given environmental variable (e.g., same osmolarity, same carbon source). This yields an MI(CDF) profile for each environmental variable across the PC spectrum, where a curve that rises steeply at low PC windows indicates that the corresponding environmental variable is encoded in the dominant modes of transcriptional variation.

Null distributions for MI were estimated by permuting environmental labels across cells while preserving within-condition transcriptional structure (per-condition shuffle), which produces a curved exponential-like null CDF. All MI calculations and spectral correlation analyses were performed using custom Python scripts within the spipy framework [Zaydman et al., 2022a, Raman et al., 2019, Gowda et al., 2022].

### Polysome profiling

Polysome profiling was performed as previously described [Aboulhouda et al., 2017] with minor modifications. Cells were grown to mid-log phase in the appropriate media, then subjected to the indicated stress conditions. Immediately prior to harvesting, cycloheximide was added to a final concentration of 100 μg/mL to stabilize polysomes. Cells were harvested by centrifugation at 3,000 × g for 3 min at 4°C, washed once with lysis buffer (20 mM Tris-HCl pH 7.4, 140 mM KCl, 5 mM MgCl_2_, 100 μg/mL cycloheximide, 1 mM DTT, 1% Triton X-100), and lysed by vortexing with glass beads. Lysates were clarified by centrifugation at 10,000 × g for 10 min at 4°C. Equal amounts of clarified lysate (by A_260_) were loaded onto 10–50% (w/v) sucrose gradients prepared in gradient buffer (20 mM Tris-HCl pH 7.4, 140 mM KCl, 5 mM MgCl_2_, 100 μg/mL cycloheximide). Gradients were centrifuged in an SW41 Ti rotor at 35,000 rpm for 2.5 h at 4°C. Fractions were collected using a Brandel gradient fractionation system with continuous monitoring of absorbance at 260 nm. Polysome-to-monosome (P/M) ratios were calculated from the integrated areas under the polysome and 80S monosome peaks.

### Laboratory evolution under selection

Directed evolution experiments were conducted using the eVOLVER system [Wong et al., 2018] in synthetic dextrose media containing 0.8 M KCl. Culture dilution was automated at OD_600_ of 0.56 and diluted to 0.08. Growth rate *k* for each growth cycle was calculated from Δ log OD ∼ *kt* (where *t* is time) based on OD ranging from 0.4 to 0.5. Evolved populations were periodically frozen and analyzed using single-cell and bulk RNA sequencing.

### Genome sequencing

Yeast genomic DNA was extracted using the acid-washed glass beads method. Whole-genome libraries were prepared using Illumina DNA Prep (Cat. # 20060059) following the manufacturer’s protocol and sequenced on a NovaSeq 6000 at the University of Chicago Genomics Facility. Adapter sequences were removed using fastp 0.23.4, reads were mapped to the *Saccharomyces cerevisiae* reference genome R64-4-1 using bwa-mem 0.7.17, and variants were identified using GATK 4.4.0 (https://github.com/broadinstitute/gatk) following GATK best practices.

### CRISPR reconstruction of derived strain

CRISPR engineering was performed as described previously [Bao et al., 2015] using the pCRCT system (Addgene plasmid # 60621), introducing the five derived-strain mutations sequentially. For each mutation, we identified potential guide RNAs near the variant site using the scoring system of Joung et al. [2017] and selected the highest-scoring gRNA. Repair templates were designed to introduce the desired mutation along with synonymous mutations at the gRNA target site to prevent repeated Cas9 cleavage. Guide RNA and repair template sequences (Table S7) were synthesized by IDT and inserted into pCRCT using the Golden Gate Assembly Kit BsaI-HF^®^ v2 (NEB, Cat. # E1601S). All engineered mutations were verified by Sanger sequencing.

### Competitive fitness assays

Growth competitions were performed by mixing an mCherry-labeled reference strain (DPY731, *TDH3pr*-mKate2) with the ancestral, derived, or CRISPR-engineered strains, respectively. Mixed cultures were grown in SDC media with or without 0.8 M KCl and diluted 1:512 daily. Cell counts from each strain were measured by flow cytometry at days 0, 1, 2, and 3.

## Acknowledgements

We thank Madhav Mani, Seppe Kuehn, Mike Rust, Ed Munro, Oni Basu, Suri Vaikuntanathan, Arvind Murugan, Allison Squires, Allan Drummond, Elizabeth Jerrison, Rama Ranganathan, and the members of the Pincus and Raman laboratories for helpful discussions and comments on the paper. We are grateful to the University of Chicago Functional Genomics Facility (RRID: SCR_019196) for scRNA-seq and bulk RNA-seq library preparation and sequencing, the University of Chicago Cytometry and Antibody Technology Facility (RRID: SCR_017760) for high-throughput flow cytometry, and the University of Chicago Integrated Light Microscopy Core (RRID: SCR_019197) for microscopy support. We set up the eVOLVER system with help from Brandon Wong and Ahmad S. Khalil. This work was supported by National Institutes of Health grants R01 GM138689 and RM1 GM153533 (to D.P.), National Science Foundation QLCI QuBBE grant OMA-2121044 (to D.P.), the Duchossois Family Institute at the University of Chicago (to A.S.R.), the Dr. Ralph and Marian Falk Medical Research Trust (to A.S.R.), NIH grant RM35 GM146702 (to A.S.R.), and the Center for Physics of Evolving Systems at the University of Chicago.

## Author contributions

**Conceptualization:** A.D., D.P., A.S.R.

**Methodology:** A.D., Y.L., B.A.D., A.A., M.G.I., L.D., V.A.

**Investigation:** A.D., Y.L., B.A.D., A.A., M.G.I., L.D., V.A.

**Formal analysis:** A.D., Y.L., B.A.D.

**Visualization:** A.D.

**Writing – original draft:** A.D., D.P., A.S.R.

**Writing – review & editing:** all authors

**Supervision:** D.P., A.S.R.

**Funding acquisition:** D.P., A.S.R.

## Data and code availability

Raw sequencing data (scRNA-seq and bulk RNA-seq) have been deposited in the Gene Expression Omnibus (GEO) under accession number GSE327265. Genome sequencing data for the derived strain have been deposited in the Sequence Read Archive (SRA) under accession number SRR37929879. Processed count matrices, metadata, and analysis notebooks are available at https://doi.org/10.5281/zenodo.19273161. Custom Python scripts for image analysis and quantification are available at the same repository. Scripts for MICDF generation are available at https://github.com/annisa-dea/complex-environments. All other data supporting the findings of this study are available from the corresponding authors upon reasonable request.

## Lead contact

Further information and requests for resources and reagents should be directed to and will be fulfilled by the lead contacts, David Pincus and Arjun S. Raman.

## Supplementary Figures

**Fig. S1:**
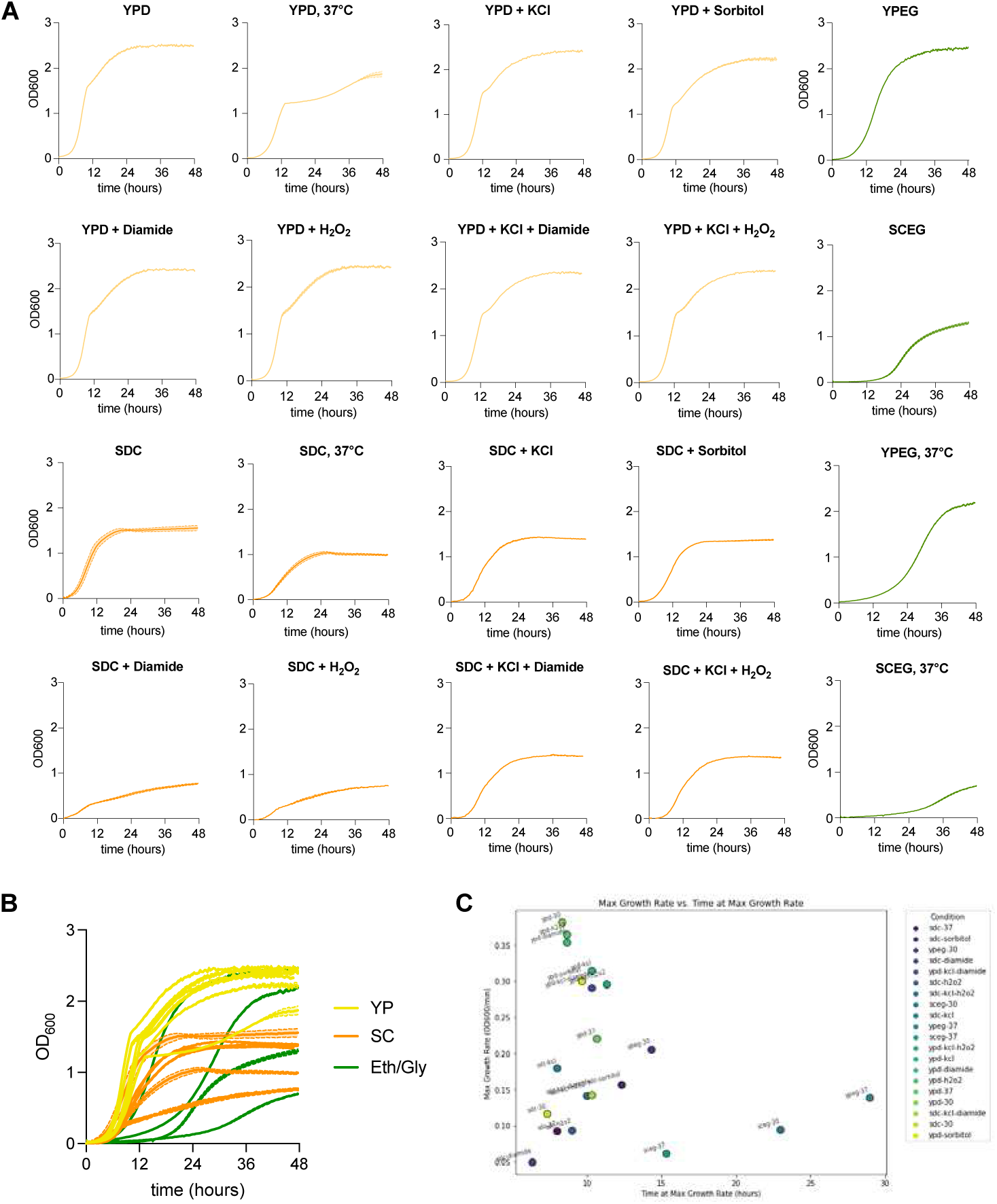
Growth curves of ancestral strain across environmental conditions. **A.** OD_600_ versus time for *S. cerevisiae* grown across 20 different complex environments. **B.** OD_600_ versus time for *S. cerevisiae* across environments shown in panel A grouped by whether environment is composed of SDC media (yellow), YPD media (orange), or ethanol/glycerol (green). **C.** Maximum growth rate across all 20 complex environments versus time to reach maximum growth rate of log phase for scRNA-seq collection.

**Fig. S2:**
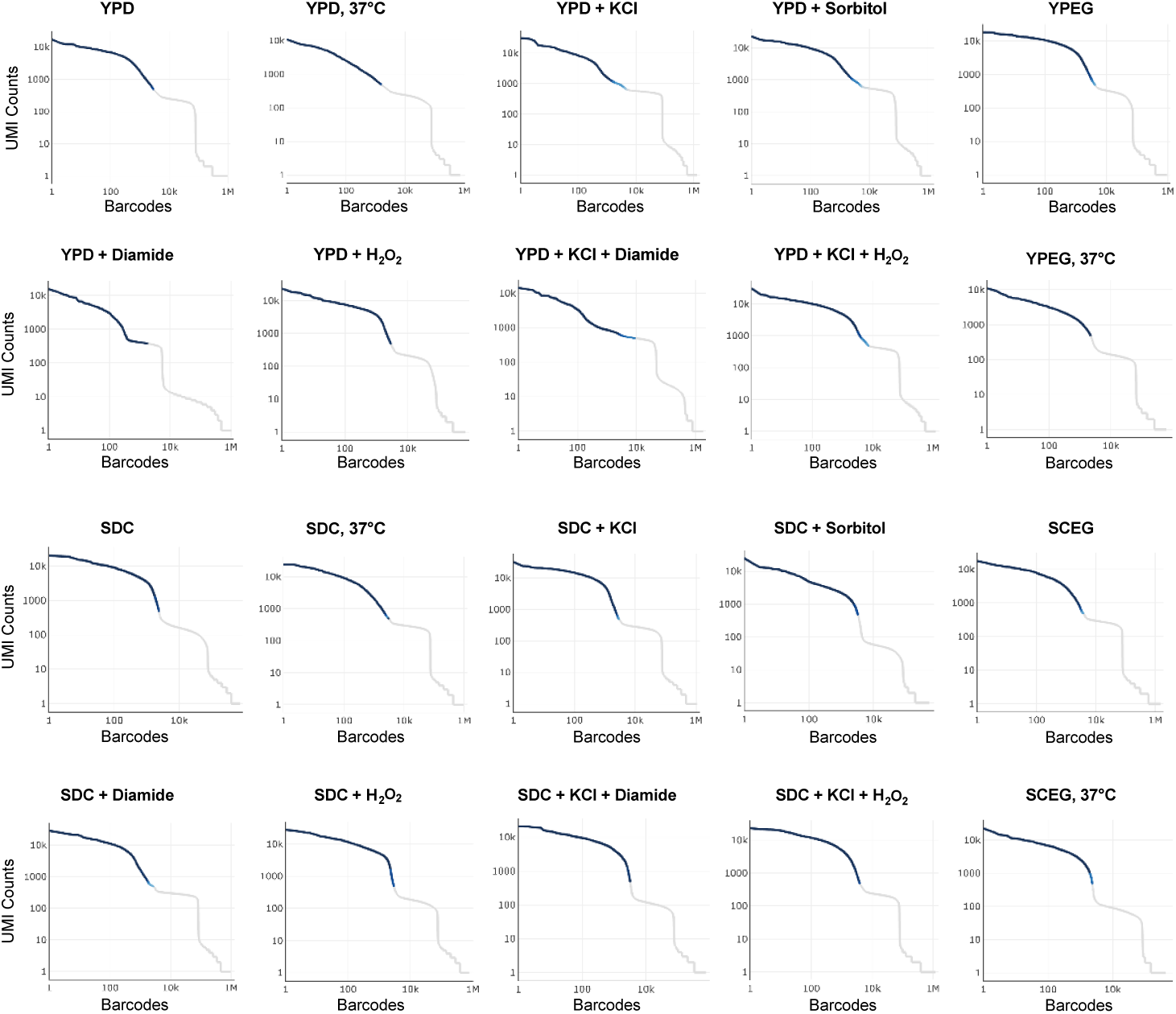
CellRanger barcode rank plots for ancestral single-cell RNA sequencing conditions. Each panel shows the UMI count per barcode ranked in decreasing order (log–log scale) for one sequencing sample. Dark blue indicates barcodes called as cells; light gray indicates empty droplets. Cell calling was performed using CellRanger v7.1.0.

**Fig. S3:**
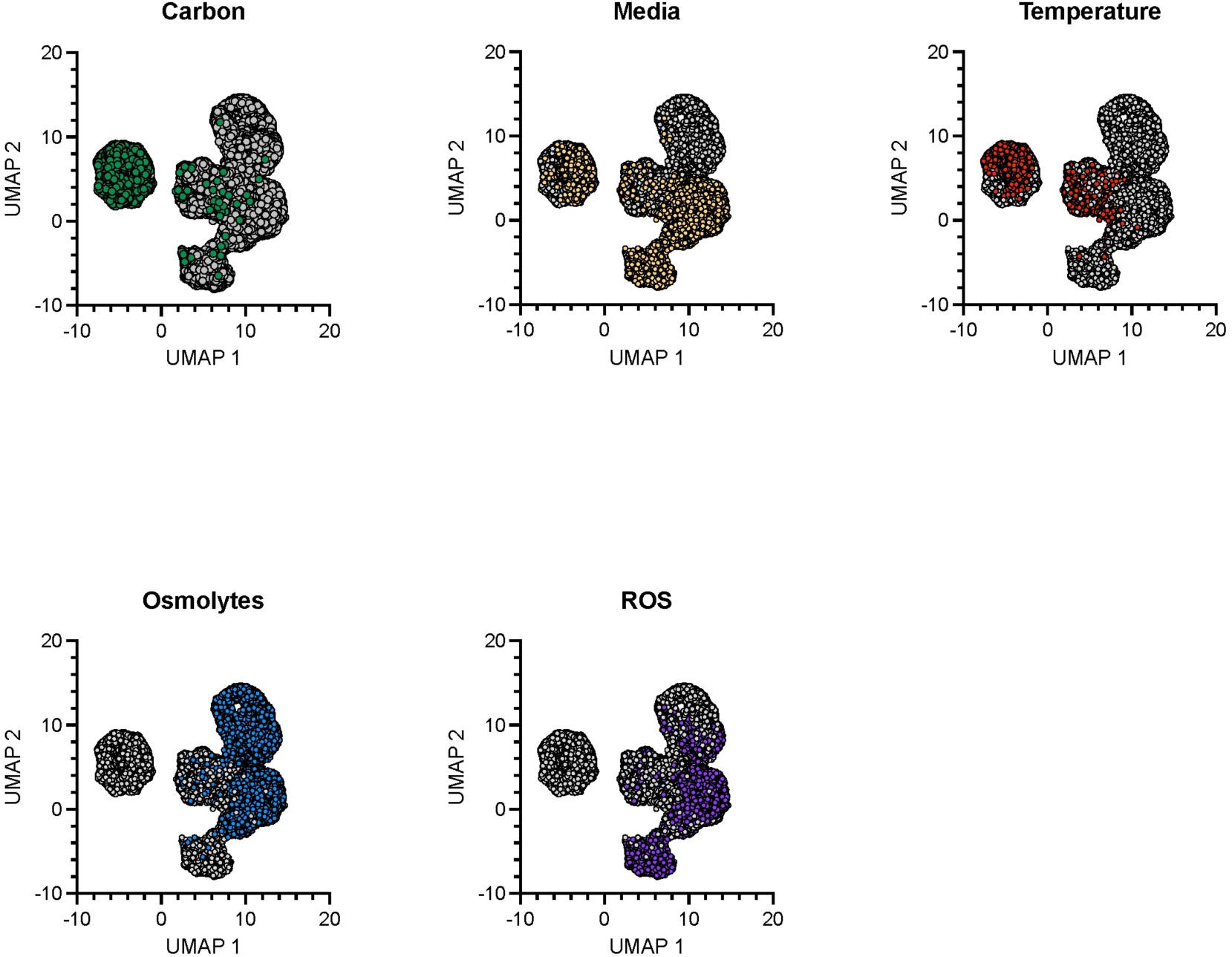
UMAP colored by individual environmental metadata. UMAP plot of cell populations shown in Fig. 1D colored by various environmental cues.

**Fig. S4:**
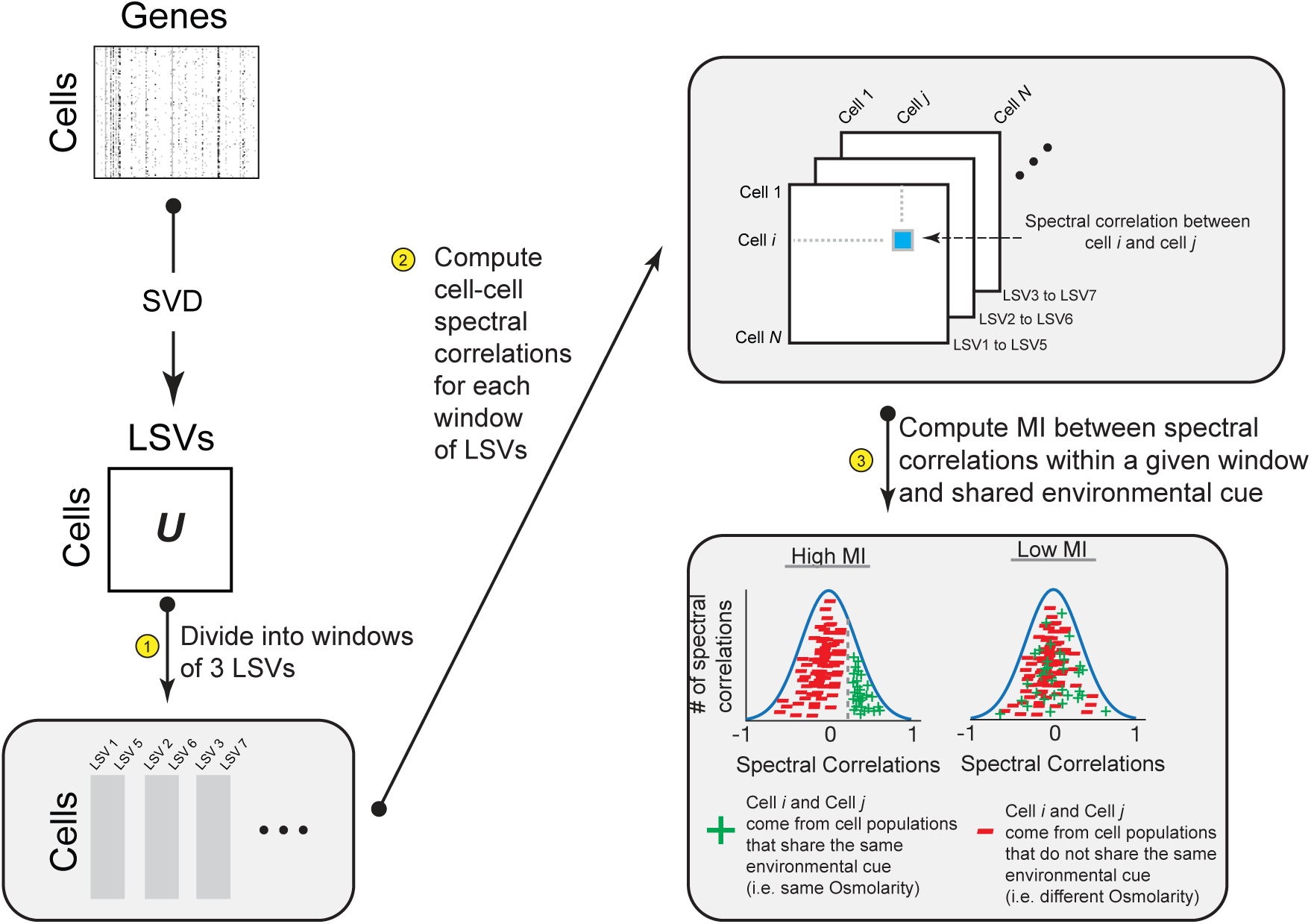
SCALES analysis of environmental information across PCs. Schematic of the Spectral Correlation Analysis of Layered Evolutionary Signals (SCALES) method. A log-normalized gene expression matrix is decomposed by singular value decomposition (SVD). SVD yields a cell embedding matrix *U* whose columns are the left singular vectors (LSVs), capturing the dominant axes of transcriptional variation across cells. (1) *U* is partitioned into overlapping windows of 3 consecutive LSVs. (2) Within each window, pairwise spectral correlations are computed between all cells, producing an *N* × *N* correlation matrix. (3) Mutual information (MI) is calculated between the distribution of spectral correlations and the shared environmental label of each cell pair. High MI indicates that the LSV window encodes transcriptional structure that distinguishes environmental conditions; low MI indicates that window carries little discriminatory information.

**Fig. S5:**
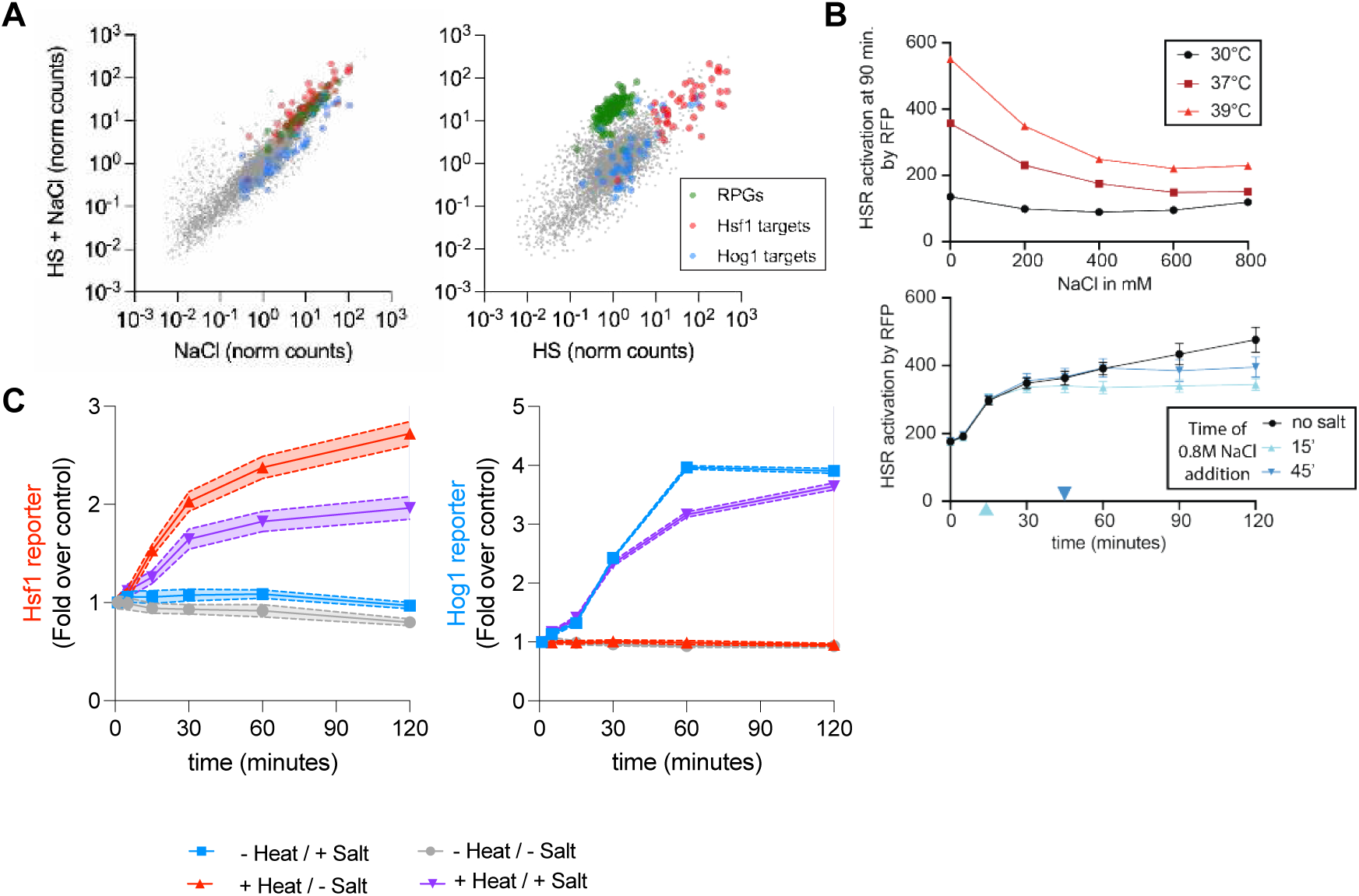
Reporter and transcriptomic validation of environmental epistasis. **A.** Transcriptome-wide comparison of gene expression in the dual stress condition (HS + NaCl) versus each individual stress. Left: dual stress versus osmotic shock (NaCl) alone; right: dual stress versus heat shock (HS) alone. Normalized read counts are shown on log–log axes with Hsf1 targets (red), Hog1 targets (blue), and ribosomal protein genes (RPGs, green) highlighted. **B.** (top) Dose response matrix of HSE-RFP fluorescent transcriptional reporter across increasing temperature (30°C, 37°C, and 39°C) and osmotic stress (0, 200, 400, 600, 800 mM of NaCl). (bottom) Heat shock time course measurements of HSE-RFP with 800 mM of NaCl introduced at 15 minutes and 45 minutes following heat shock. Each data point represents mean HSE-RFP fluorescence normalized to side scatter (SSC) across 3 biological replicates. **C.** Transcriptional fluorescence reporter measurements of HSE-RFP and pHor2-GFP of cells grown in 2% ethanol/2% glycerol media. Cells are subjected to heat shock (39°C), osmotic shock (0.8 M NaCl), or simultaneous dual stress. Stress response measurements are lower in ethanol/glycerol-grown media compared to glucose media (Fig. 2E).

**Fig. S6:**
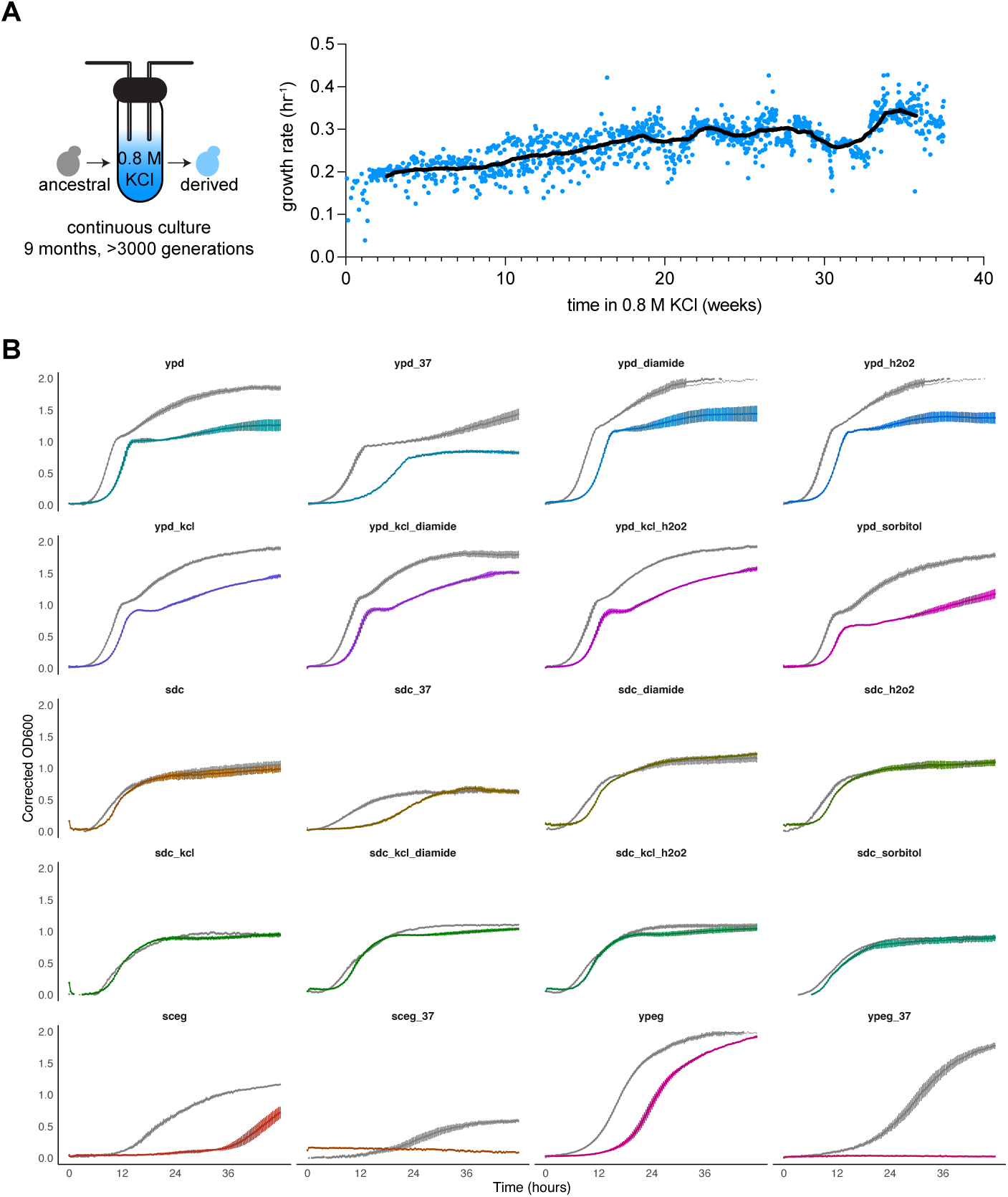
Growth characteristics of the derived strain. **A.** Growth rate of the evolving strain versus number of weeks of selection under 0.8 M KCl. **B.** OD_600_ growth curve measurements versus time of the derived strain (color), plotted against the ancestral strain (gray). The derived strain exhibited slower growth in most conditions, with the exception of the selection condition. The derived strain is unable to grow at 37°C under ethanol/glycerol media.

**Fig. S7:**
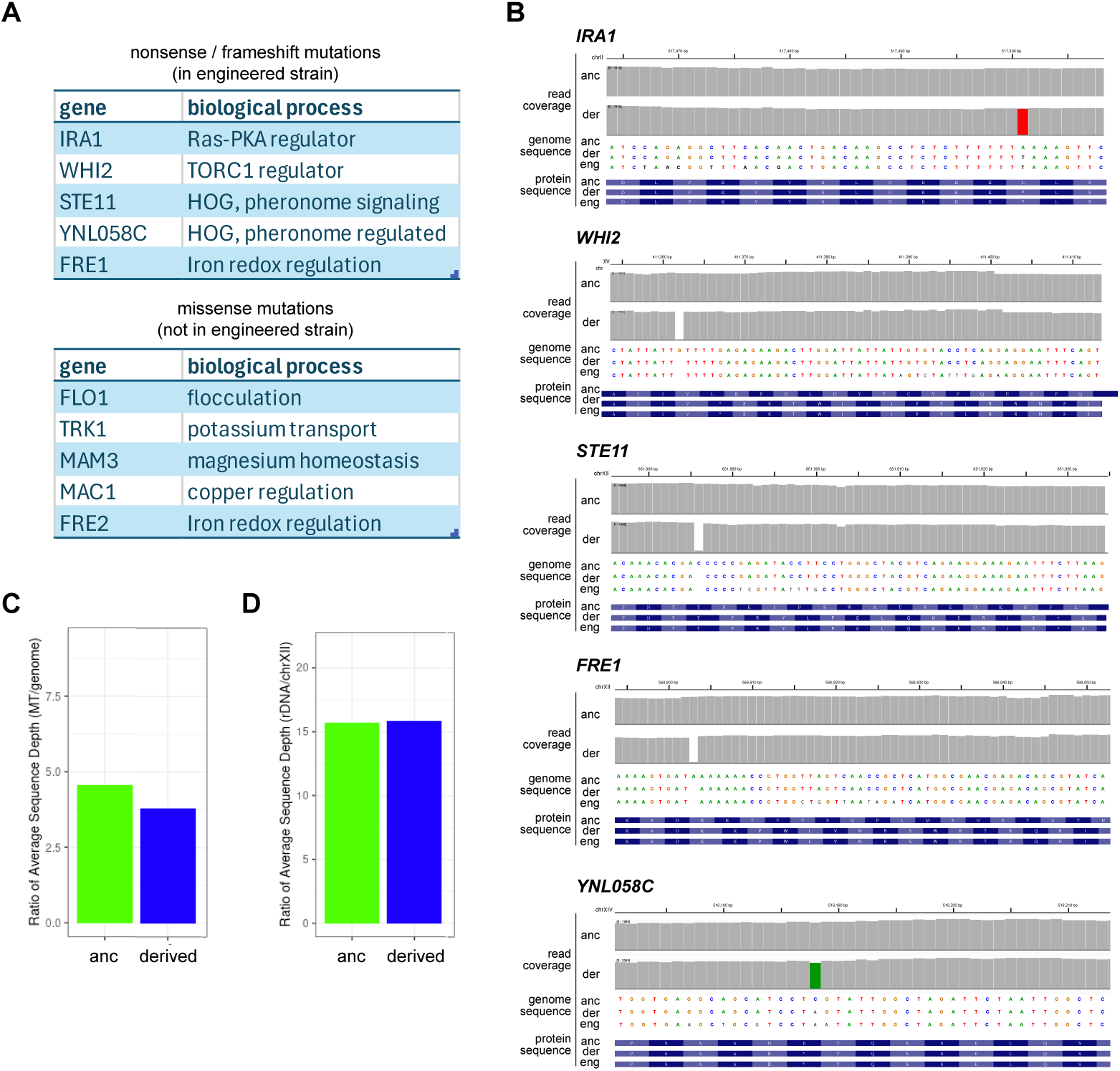
Genomic characterization of the derived strain. **A.** Genome-wide view of mutations in the derived strain relative to the ancestral strain, showing nonsense and frameshift mutations (top) and missense mutations (bottom) across all chromosomes. **B.** Read coverage and sequence alignments at the five genes harboring premature termination codons (*IRA1*, *WHI2*, *STE11*, *FRE1*, and *YNL058C*), comparing the parental (P), derived (V0), and CRISPR-engineered (5gm) strains. **C.** Ratio of average sequencing depth for the mitochondrial genome relative to the nuclear genome in ancestral and derived strains. **D.** Ratio of average sequencing depth for the rDNA locus on chromosome XII relative to the rest of the nuclear genome in ancestral and derived strains.

**Fig. S8:**
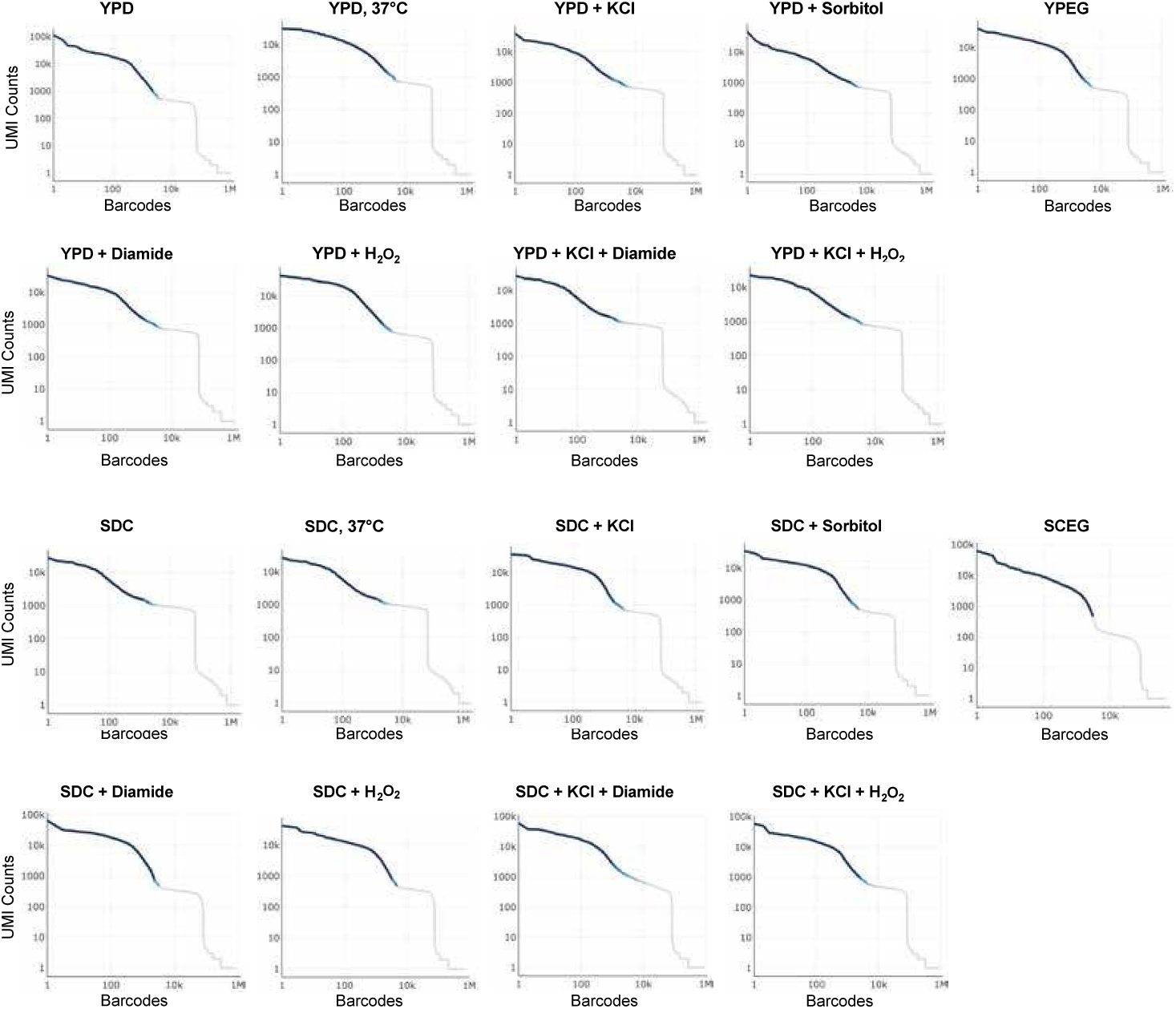
CellRanger barcode rank plots for derived strain single-cell RNA sequencing conditions. Each panel shows the UMI count per barcode ranked in decreasing order (log–log scale) for one sequencing sample. Dark blue indicates barcodes called as cells; light gray indicates empty droplets. Cell calling was performed using CellRanger v7.1.0.

**Fig. S9:**
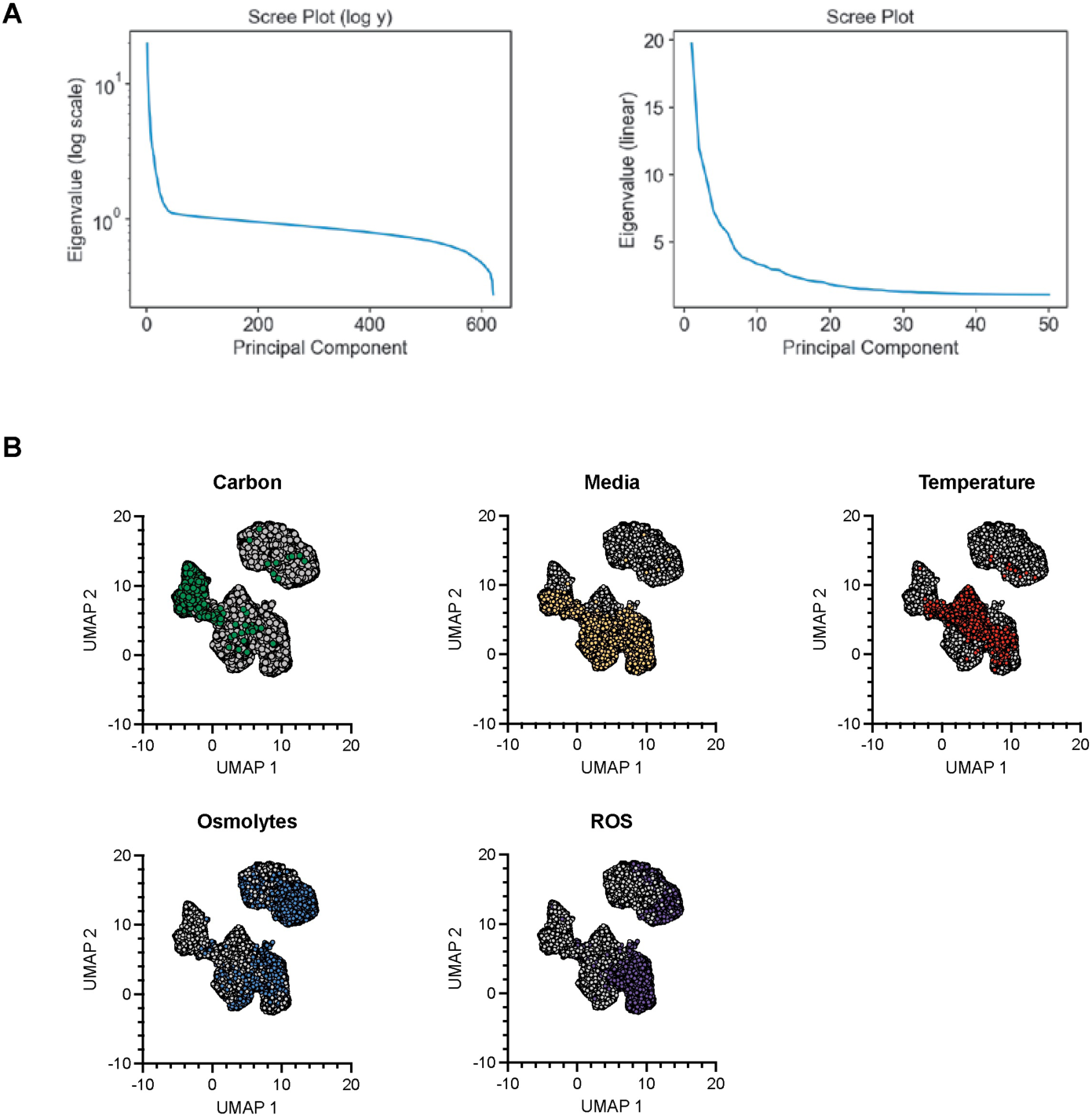
Scree plots and UMAP analysis of the derived strain scRNA-seq data. **A.** Fraction of total variance explained by each left singular vector (LSV) following SVD of the log-normalized gene expression matrix of the derived strain. **B.** UMAP plot of derived strain cell populations colored by various environmental cues.

**Fig. S10:**
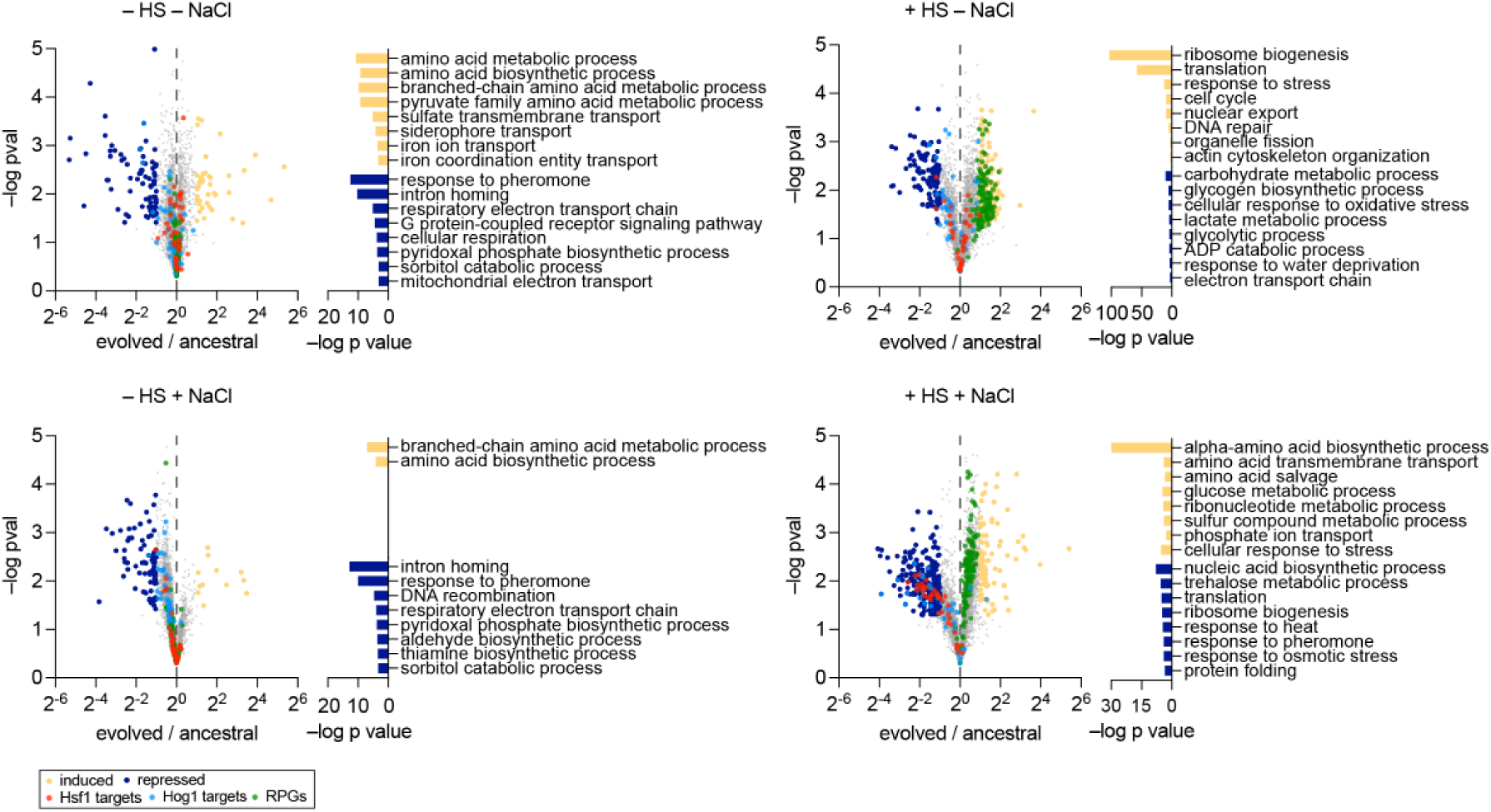
Differential gene expression and GO term enrichment in derived cells across stress conditions. Differential gene expression between the derived and ancestral strains across four conditions: unstressed (−HS −NaCl), heat shock (+HS −NaCl), osmotic shock (−HS +NaCl), and dual stress (+HS +NaCl). (Left on each panel) Volcano plots showing bulk RNA-seq differential expression between derived and ancestral cells. Colored points indicate Hsf1 target genes (red), Hog1 target genes (green), and ribosomal protein genes (RPGs, blue). (Right on each panel) GO biological process enrichment of genes induced (orange) or repressed (blue) in evolved cells.

## Supplementary Tables

**Table S1:** QC metrics of single-cell sequencing runs. Summary statistics from CellRanger v7.1.0 for all ancestral (anc) and evolved (evo) scRNA-seq samples.

| Condition | Reads | Valid BC (%) | Sat. (%) | Q30 (%) | Map Genome (%) | Map Txome (%) | Cells | Med. Genes | Med. UMI |
| --- | --- | --- | --- | --- | --- | --- | --- | --- | --- |
| *Ancestral strain* |  |  |  |  |  |  |  |  |  |
| SCEG-30 | 160M | 97.3 | 66.0 | 95.6 | 75.4 | 53.5 | 3,353 | 902 | 1,688 |
| SCEG-37 | 160M | 96.7 | 80.9 | 96.0 | 64.1 | 39.9 | 2,212 | 1,172 | 2,416 |
| SDC | 145M | 97.4 | 79.5 | 94.8 | 78.8 | 60.1 | 2,313 | 1,219 | 3,119 |
| SDC-37 | 172M | 97.3 | 70.0 | 95.5 | 75.4 | 52.4 | 2,767 | 902 | 1,429 |
| SDC-diamide | 146M | 97.4 | 59.4 | 95.7 | 74.4 | 51.7 | 2,056 | 844 | 1,620 |
| SDC-H<sub>2</sub>O<sub>2</sub> | 158M | 97.4 | 69.4 | 95.2 | 76.3 | 51.9 | 2,813 | 1,362 | 4,314 |
| SDC-KCl | 169M | 98.0 | 72.2 | 94.8 | 85.2 | 67.3 | 2,756 | 1,348 | 3,271 |
| SDC-KCl-dia | 147M | 97.7 | 77.5 | 95.4 | 82.9 | 59.4 | 3,191 | 1,303 | 2,958 |
| SDC-KCl-H<sub>2</sub>O<sub>2</sub> | 175M | 97.4 | 69.7 | 95.0 | 79.5 | 57.7 | 3,844 | 1,206 | 2,644 |
| SDC-sorbitol | 132M | 97.6 | 85.4 | 95.7 | 78.5 | 54.3 | 3,123 | 939 | 1,701 |
| YPD | 149M | 97.1 | 73.6 | 95.8 | 75.9 | 57.7 | 2,844 | 744 | 1,180 |
| YPD-37 | 142M | 96.5 | 74.7 | 93.3 | 68.5 | 46.7 | 1,534 | 522 | 828 |
| YPD-diamide | 140M | 97.7 | 93.5 | 95.4 | 82.2 | 62.2 | 1,896 | 311 | 416 |
| YPD-H<sub>2</sub>O<sub>2</sub> | 164M | 97.1 | 79.1 | 95.7 | 76.3 | 57.8 | 2,862 | 1,217 | 2,506 |
| YPD-KCl | 159M | 97.9 | 46.1 | 95.8 | 81.8 | 59.5 | 2,846 | 653 | 1,262 |
| YPD-KCl-dia | 142M | 97.3 | 61.6 | 94.6 | 79.6 | 56.6 | 6,343 | 358 | 590 |
| YPD-KCl-H<sub>2</sub>O<sub>2</sub> | 163M | 97.8 | 54.0 | 95.7 | 85.4 | 60.7 | 5,440 | 960 | 1,968 |
| YPD-sorbitol | 130M | 97.6 | 41.0 | 95.6 | 81.7 | 60.3 | 4,236 | 680 | 1,286 |
| YPEG-30 | 160M | 97.7 | 67.4 | 95.1 | 81.0 | 58.3 | 3,659 | 1,078 | 2,178 |
| YPEG-37 | 115M | 96.5 | 75.6 | 95.3 | 65.7 | 40.9 | 2,260 | 675 | 1,113 |
| *Derived strain* |  |  |  |  |  |  |  |  |  |
| SCEG-30 | 138M | 95.8 | 75.8 | 96.8 | 73.1 | 52.6 | 3,081 | 1,008 | 2,092 |
| SDC | 159M | 97.7 | 61.1 | 96.9 | 84.0 | 61.8 | 3,114 | 1,106 | 2,368 |
| SDC-37 | 166M | 98.2 | 55.1 | 97.0 | 87.3 | 66.4 | 3,623 | 1,181 | 2,667 |
| SDC-diamide | 168M | 96.9 | 54.3 | 97.0 | 80.6 | 50.4 | 2,663 | 1,108 | 2,578 |
| SDC-H<sub>2</sub>O<sub>2</sub> | 158M | 97.9 | 58.2 | 97.0 | 88.6 | 61.4 | 4,040 | 1,061 | 2,030 |
| SDC-KCl | 177M | 98.1 | 48.0 | 97.0 | 89.5 | 59.8 | 3,571 | 783 | 1,607 |
| SDC-KCl-dia | 164M | 98.2 | 52.8 | 97.0 | 89.4 | 64.8 | 3,421 | 670 | 1,312 |
| SDC-KCl-H<sub>2</sub>O<sub>2</sub> | 141M | 97.3 | 49.6 | 96.8 | 82.1 | 65.1 | 3,573 | 708 | 1,379 |
| SDC-sorbitol | 149M | 97.0 | 53.0 | 97.0 | 79.4 | 57.2 | 3,792 | 832 | 1,426 |
| YPD | 137M | 96.4 | 50.1 | 96.9 | 78.4 | 61.9 | 2,997 | 1,011 | 1,924 |
| YPD-37 | 179M | 98.2 | 45.4 | 97.0 | 88.3 | 59.3 | 4,127 | 863 | 1,705 |
| YPD-diamide | 141M | 98.2 | 43.7 | 96.6 | 87.2 | 69.8 | 2,842 | 670 | 1,367 |
| YPD-H<sub>2</sub>O<sub>2</sub> | 157M | 97.4 | 47.4 | 97.0 | 81.8 | 62.3 | 3,090 | 718 | 1,610 |
| YPD-KCl | 140M | 97.2 | 38.2 | 97.0 | 82.7 | 66.8 | 4,165 | 570 | 1,084 |
| YPD-KCl-dia | 167M | 98.4 | 44.8 | 97.1 | 89.1 | 70.7 | 2,747 | 594 | 1,508 |
| YPD-KCl-H<sub>2</sub>O<sub>2</sub> | 147M | 97.4 | 42.5 | 96.8 | 82.2 | 65.8 | 3,254 | 598 | 1,322 |
| YPD-sorbitol | 118M | 96.7 | 36.9 | 96.9 | 78.6 | 62.1 | 4,773 | 637 | 1,084 |
| YPEG-30 | 171M | 98.0 | 61.4 | 97.0 | 88.5 | 66.9 | 3,779 | 799 | 1,518 |

**Table S2:**
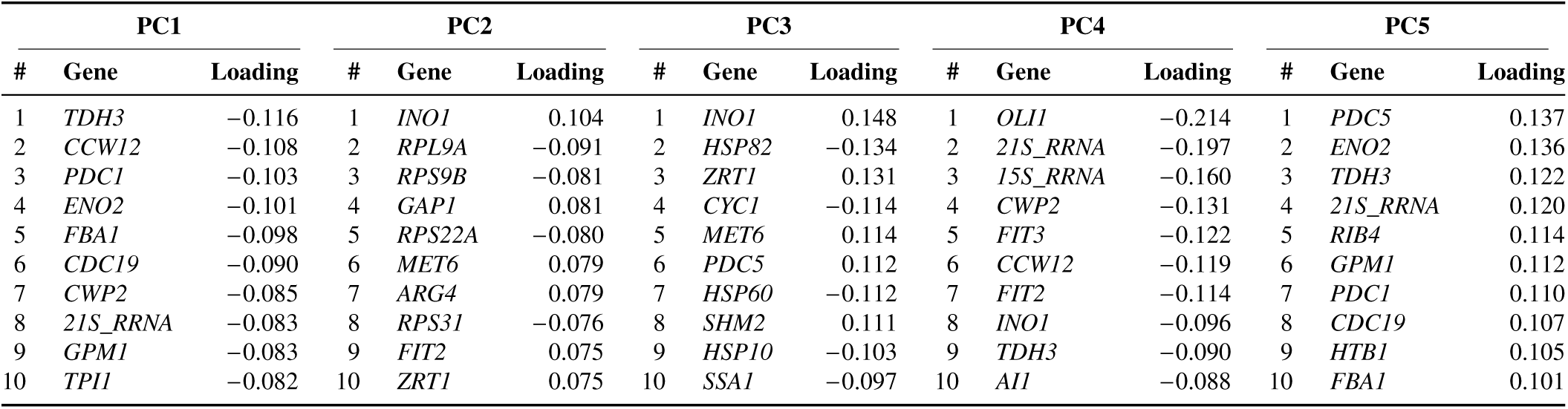
Top 10 gene loadings per principal component (ancestral strain). For each of the first five principal components from SVD of the ancestral scRNA-seq dataset, the 10 genes with the largest absolute loadings are shown with their loading values. Positive and negative loadings indicate opposing directions of transcriptional variation along each PC.

**Table S3:**
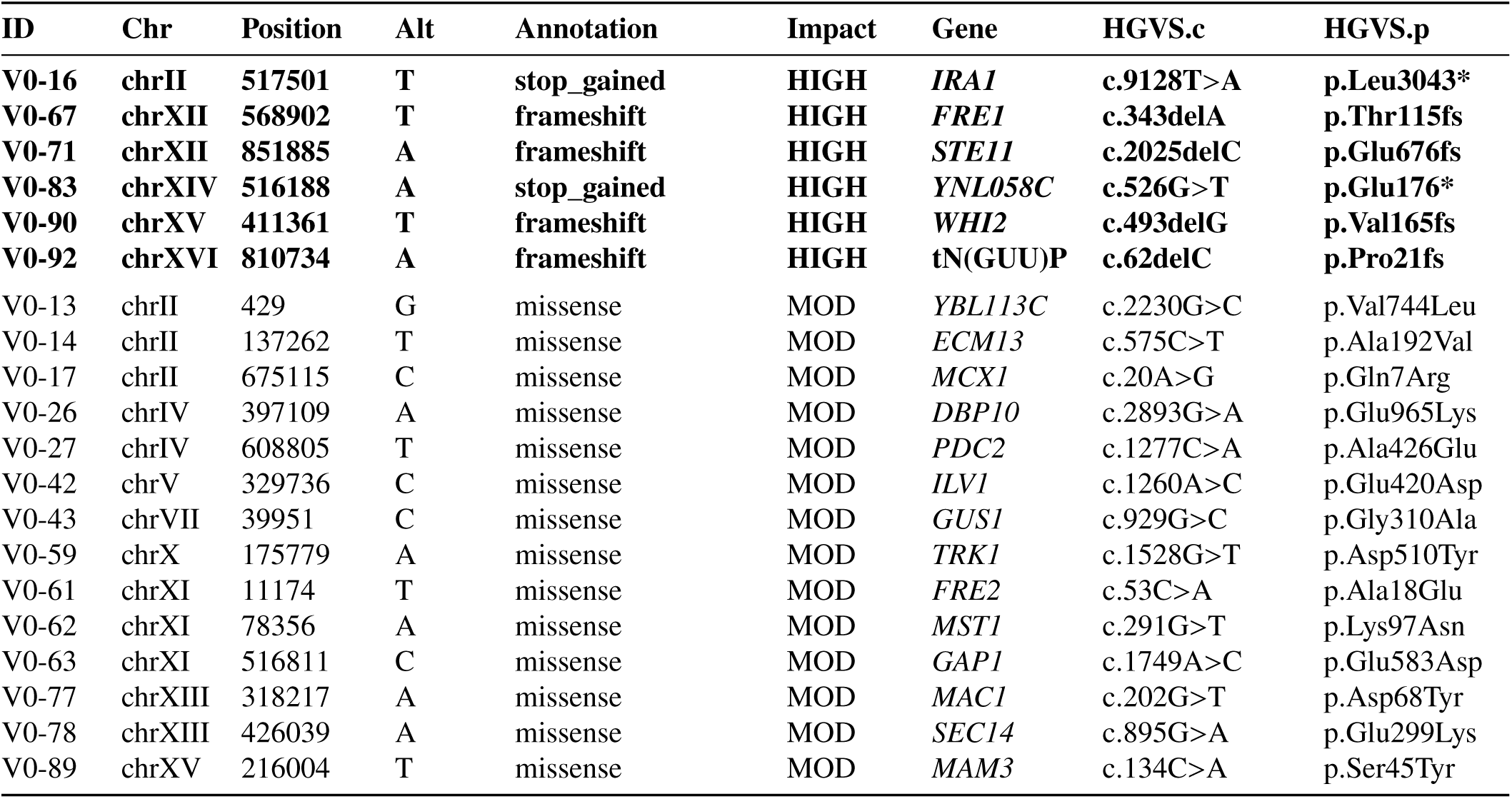
Genomic variants in the derived strain (HIGH and MODERATE impact). Variants were identified by whole-genome sequencing and annotated using SnpEff relative to the *S. cerevisiae* R64-4-1 reference genome. Only variants with HIGH or MODERATE (MOD) predicted impact are shown. HIGH-impact variants (premature stop codons and frameshifts) are shown in bold.

**Table S4:**
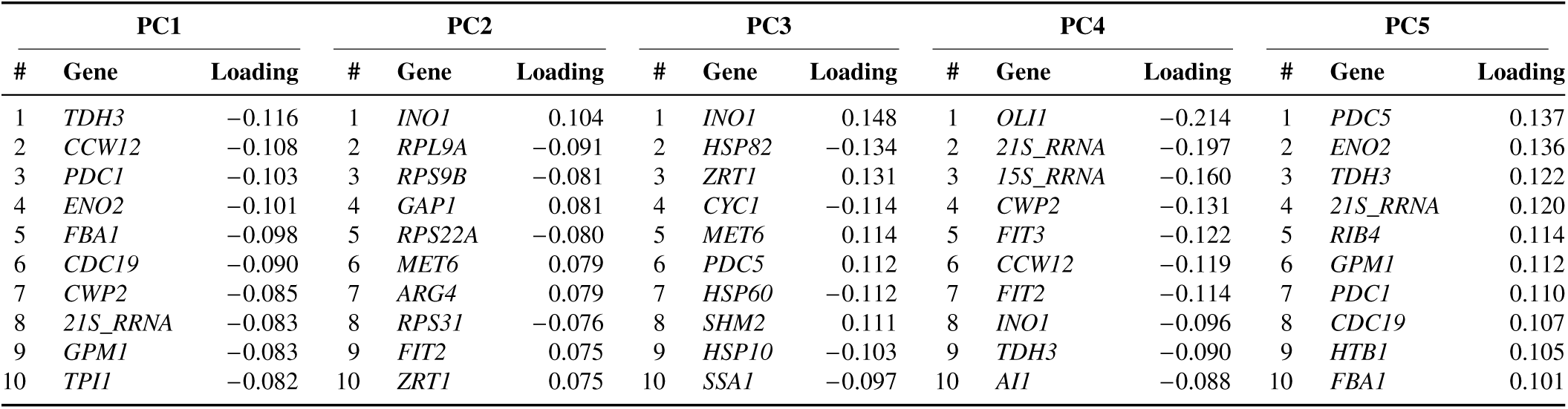
Top 10 gene loadings per principal component (derived strain). For each of the first five principal components from SVD of the derived scRNA-seq dataset, the 10 genes with the largest absolute loadings are shown with their loading values. Positive and negative loadings indicate opposing directions of transcriptional variation along each PC.

**Table S5:** Yeast strains used in this study. All strains are in the W303 background (*MAT**a***; *leu2-3,112; trp1-1; can1-100; ura3-1; ade2-1; his3-11,15*).

| Strain | Genotype | Bg | Description | Figures |
| --- | --- | --- | --- | --- |
| DPY144 | 4×HSE-Venus::LEU2 | W303 | WT HSE reporter | S1, 2D, 2F, 3C, 3E, 3F, S3D, 5B, S5 |
| DPY1284 | *tpk1/2/3-as*; *TOR1-1*(S1972I); *fpr1Δ*::NAT; TEF2pr-mKate-URA3-4×HSEve-EmGFP; RPL13A-2×FKBP12::TRP; SIS1-FRB::HIS | W303 | Sis1 anchor-away | 1E, 2B |
| DPY1368 | HSEv2-Venus::LEU2; *ssa1Δ*::HYG; *ssa2Δ*::KAN; TEF1pr-SSA2::HIS; *ssa4Δ*::NAT; TEF1pr-SSA2::URA; *ssa3Δ*::BLE | W303 | ΔFBL Hsp70 | 3C, 3E, 3F, S3C, 5A, 5B |
| DPY1600 | 4×HSE-Venus::LEU2; Hsp82-ERVB1-mScarlet::KAN | W303 | Hsp82-P2A-mScarlet | S1, S3C, 5A |
| DPY1615 | 4×HSE-Venus::LEU2; Sis1-ERVB1-mScarlet::KAN | W303 | SIS1pr-SIS1-P2A-mScarlet | S3A, S3B, S3D |
| DPY1661 | 4×HSE-Venus::LEU2; Sup35pr-Sis1-ERVB1-mScarlet::KAN | W303 | Sup35pr-SIS1-P2A-mScarlet | S3A, S3B, S3D |
| DPY1701 | 4×HSE-Venus::LEU2; Sup35pr-SIS1 | W303 | Sup35pr-SIS1 | S3C, S3D |
| DPY1761 | 4×HSE-Venus::LEU2; Sup35pr-SIS1; Sup35pr-Sis1::TRP | W303 | 2× Sup35pr-SIS1 | 3C, 3E, 3F, S3C, S3D, 5B, 5A, S5 |
| DPY1846 | Sis1-mVenus::HIS; Nsr1-mScarletI::KAN; *ssa1pr-HALO-ssa1ORF*::LEU | W303 | Ssa1-HaloTag imaging | 4B, 4D |
| DPY1930 | HALO-Sis1; Nsr1-mScarletI::KAN | W303 | Sis1-HaloTag imaging | 4B, 4C |
| DPY1990 | Hsf1-mVenus::HIS; *ssa1pr-HALO-ssa1ORF*::LEU | W303 | Ssa1-HaloTag; Hsf1-mVenus | 4E, 4F |
| DPY1991 | 4×HSE-mVenus::HIS; Pab1-mKate::URA3 | W303 | Pab1-mKate | 5C |
| DPY1992 | 4×HSE-Venus::LEU2; Sup35pr-SIS1; Sup35pr-Sis1::TRP; Pab1-mKate::URA3 | W303 | Pab1-mKate; 2× Sup35pr-SIS1 | 5C |
| DPY2001 | HALO-Sis1 (N-terminal CRISPR); Hsf1-mVenus::HIS | W303 | HALO-Sis1; Hsf1-mVenus | 4E, 4F |
| *Additional strains* |  |  |  |  |
| — | LEU2::HSE-mApple; TRP1::pHOR2-GFP | W303 | HSR/HOG dual reporter | 2 |
| — | Hog1-[FP]; Hsf1-[FP] (endogenous tagged) | W303 | Hog1/Hsf1 nuclear localization | 3B, 3C |
| — | Sis1-Venus::HIS3; Nsr1-mScarletI::KAN; Rpl26a-HaloTag | W303 | oRP/Sis1 co-localization | 3E, 3F |
| Derived (V0) | W303 + IRA1(L3043*); STE11(E676fs); WHI2(V165fs); FRE1(T115fs); YNL058C(E176*); petite | W303<sup>†</sup> | Derived strain (>3000 gen. KCl) | 4–6, S6–S8 |
| 5gm (CRISPR) | W303 + IRA1(L3043*); STE11(E676fs); WHI2(V165fs); FRE1(T115fs); YNL058C(E176*) | W303 | CRISPR reconstruction | 4B, S7B |
| DPY731 | TDH3pr-mKate2 | W303 | mKate2 reference strain | 4B |

**Table S6:** Environmental conditions. The 20 environments used for transcriptional profiling and growth assays. YP = yeast extract + peptone (rich); SC = synthetic complete (defined). All conditions at 30°C unless otherwise noted.

| # | Label | Media | Carbon source | Temp. | Osmolyte | ROS agent | Notes |
| --- | --- | --- | --- | --- | --- | --- | --- |
| 1 | YPD-30 | YP | 2% glucose | 30°C | — | — | Reference |
| 2 | YPD-37 | YP | 2% glucose | 37°C | — | — | Heat stress |
| 3 | YPD-KCl | YP | 2% glucose | 30°C | 500 mM KCl | — | Ionic osmotic |
| 4 | YPD-sorbitol | YP | 2% glucose | 30°C | 500 mM sorbitol | — | Non-ionic osmotic |
| 5 | YPD-diamide | YP | 2% glucose | 30°C | — | 50 μM diamide | Thiol oxidant |
| 6 | YPD-H<sub>2</sub>O<sub>2</sub> | YP | 2% glucose | 30°C | — | 50 μM H<sub>2</sub>O<sub>2</sub> | ROS |
| 7 | YPD-KCl-diamide | YP | 2% glucose | 30°C | 500 mM KCl | 50 μM diamide | Osmotic + ROS |
| 8 | YPD-KCl-H<sub>2</sub>O<sub>2</sub> | YP | 2% glucose | 30°C | 500 mM KCl | 50 μM H<sub>2</sub>O<sub>2</sub> | Osmotic + ROS |
| 9 | SDC-30 | SC | 2% glucose | 30°C | — | — | Defined media |
| 10 | SDC-37 | SC | 2% glucose | 37°C | — | — | Defined + heat |
| 11 | SDC-KCl | SC | 2% glucose | 30°C | 500 mM KCl | — | Evolution condition* |
| 12 | SDC-sorbitol | SC | 2% glucose | 30°C | 500 mM sorbitol | — | Defined + non-ionic |
| 13 | SDC-diamide | SC | 2% glucose | 30°C | — | 50 μM diamide | Defined + thiol oxidant |
| 14 | SDC-H<sub>2</sub>O<sub>2</sub> | SC | 2% glucose | 30°C | — | 50 μM H<sub>2</sub>O<sub>2</sub> | Defined + ROS |
| 15 | SDC-KCl-diamide | SC | 2% glucose | 30°C | 500 mM KCl | 50 μM diamide | Defined + osmotic/ROS |
| 16 | SDC-KCl-H<sub>2</sub>O<sub>2</sub> | SC | 2% glucose | 30°C | 500 mM KCl | 50 μM H<sub>2</sub>O<sub>2</sub> | Defined + osmotic/ROS |
| 17 | YPEG-30 | YP | 2% EtOH/2% glycerol | 30°C | — | — | Non-fermentable |
| 18 | YPEG-37 | YP | 2% EtOH/2% glycerol | 37°C | — | — | Non-ferm. + heat |
| 19 | SCEG-30 | SC | 2% EtOH/2% glycerol | 30°C | — | — | Defined + non-ferm. |
| 20 | SCEG-37 | SC | 2% EtOH/2% glycerol | 37°C | — | — | Defined + non-ferm. + heat<sup>†</sup> |

\*eVOLVER evolution used 0.8 M KCl in SDC. <sup>†</sup>Derived strain failed to grow in this condition.

**Table S7:**
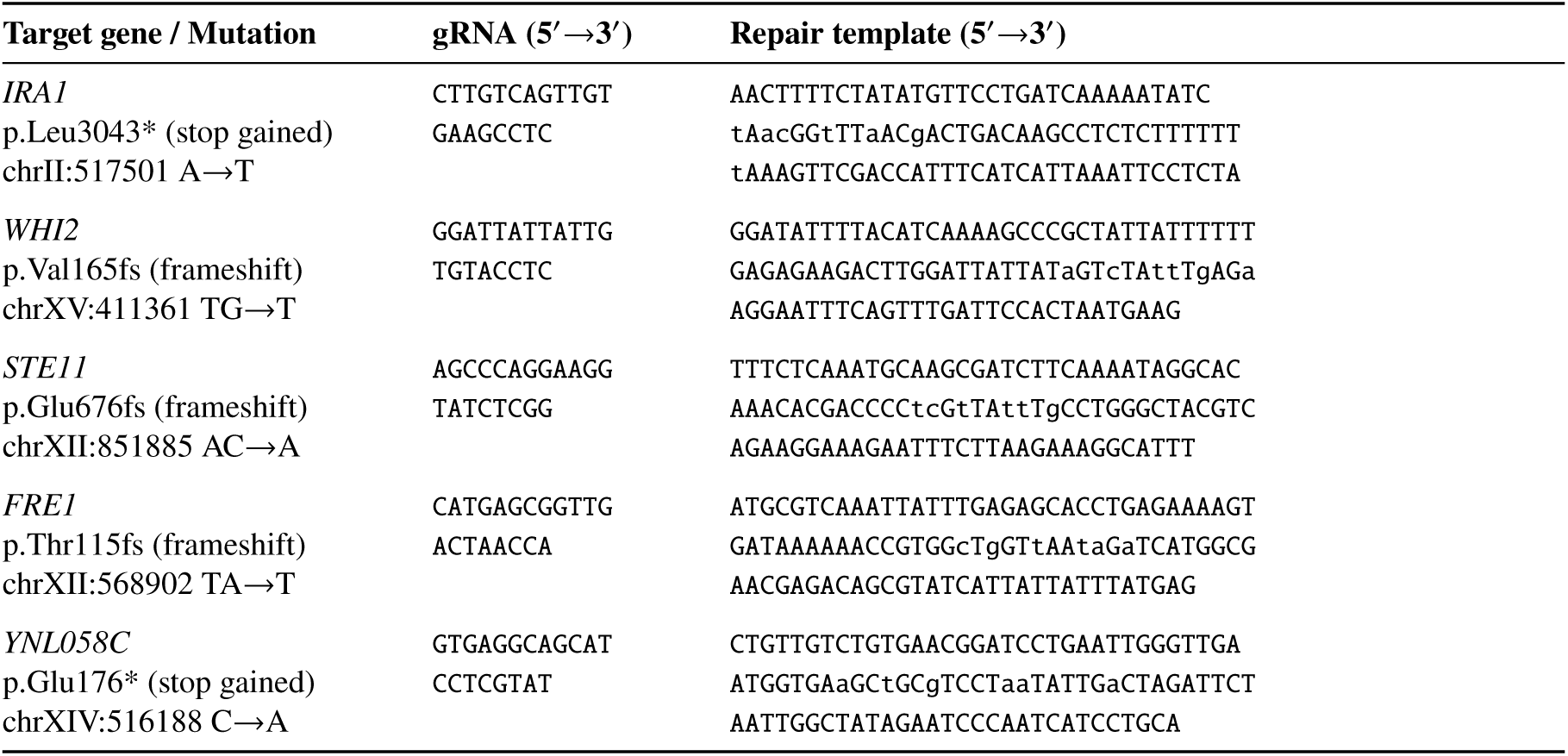
CRISPR guide RNA and repair template sequences. Guide RNAs were designed near each variant site and scored using the system of Joung et al. [2017]. Repair templates introduce the derived-strain mutation along with synonymous PAM-site mutations (lowercase) to prevent repeated Cas9 cleavage. Engineering was performed using the pCRCT system [Bao et al., 2015] (Addgene #60621).

